# Signaling input from the polar flagellum uncouples quorum sensing from cell-density regulation in *Vibrio* species

**DOI:** 10.64898/2026.06.17.732909

**Authors:** Sreeja Shaw, Sandra Sanchez, Anjali Steenhaut, Karlen Enid Correa Velez, Pantong Wu, Julia C. van Kessel, Wai-Leung Ng

**Affiliations:** Department of Molecular Biology and Microbiology, Tufts University School of Medicine, Boston, MA 02111, USA; Department of Biology, Indiana University-Bloomington, Bloomington, IN 47405, USA; Department of Biology, Framingham State University, Framingham, MA 01701, USA; Department of Biology, Tufts University, Medford, MA 02155, USA

## Abstract

Quorum sensing (QS) is traditionally recognized as a signaling mechanism that monitors cell density to coordinate bacterial group behaviors. Although the architecture of many QS pathways and their dependence on cell density are well established, the impact from other physiological cues on QS response is largely unclear.

We demonstrate here that flagellar integrity profoundly influences QS responses across multiple *Vibrio* species. Systematic disruption of flagellar components in *Vibrio cholerae* revealed that loss of the hook cap protein FlgD induces QS sRNAs Qrr1–4 production independent of cell density and the canonical QS regulator LuxO. This bypass mechanism also overrides normal cell density-dependent regulation of several QS-controlled phenotypes. Notably, Δ*flgD* mutation restores intestinal colonization of Δ*luxO* mutants in an infant mouse model, indicating that flagellar defects reshape *V. cholerae* QS during infection. This bypass mechanism is evolutionarily conserved as Δ*flgD* mutation induced LuxO-independent *qrr* expression in multiple *Vibrio* species.

The FlrBC two-component system, essential for flagellar gene expression, is required for the activation of *qrr* expression in Δ*flgD* mutants. FlgD specifically interacts with the FlrB histidine kinase but not the FlrC response regulator, and a hyperactive FlrC variant is sufficient to drive *qrr* expression even in cells with FlgD. These findings suggest that not only FlrBC senses cytoplasmic FlgD levels to monitor flagellar completeness to direct flagellar gene expression, this signaling system also functions as a link to bypass the canonical cell-density control of QS by integrating cellular structural information to coordinate group behaviors.

## INTRODUCTION

Quorum Sensing (**QS**) is a cell-cell communication process that relies on the production, detection, and response to extracellular chemical signals called autoinducers [1–3]. QS allows bacteria to monitor cell density to control population-wide gene expression and function as coordinated groups in complex communities. *Vibrio cholerae*, the human pathogen that causes the diarrheal disease cholera, uses QS to regulate many important processes in its life cycle including virulence, biofilm formation, metabolic homeostasis, genetic competence, Type VI secretion, and phage defense [4–21], underscoring the fundamental importance of microbial communication in shaping bacterial physiology, community behavior, and pathogenesis.

We previously established that four histidine kinases, CqsS, LuxPQ, CqsR, and VpsS, function in parallel to control QS in *V. cholerae* [13, 14] (**Fig. 1A**). Central to this QS circuit is the phosphorylation level of the response regulator LuxO and the total amount of the four small RNAs Qrr1-4, because they dictate the QS response of the pathogen [4, 5, 13, 14, 22]. At low cell density (**LCD**), when the extracellular autoinducer concentrations are low, the four QS histidine kinases phosphorylate LuxO via the phosphotransfer protein LuxU. Phosphorylated LuxO activates the transcription of Qrr1-4 sRNAs, which promote and inhibit the translation of transcriptional regulators AphA and HapR, respectively [22, 23]. Because AphA is an activator and HapR is a repressor of biofilm and virulence genes, these genes are activated at LCD [24–29] (**Fig. 1A**).

**Fig. 1.**
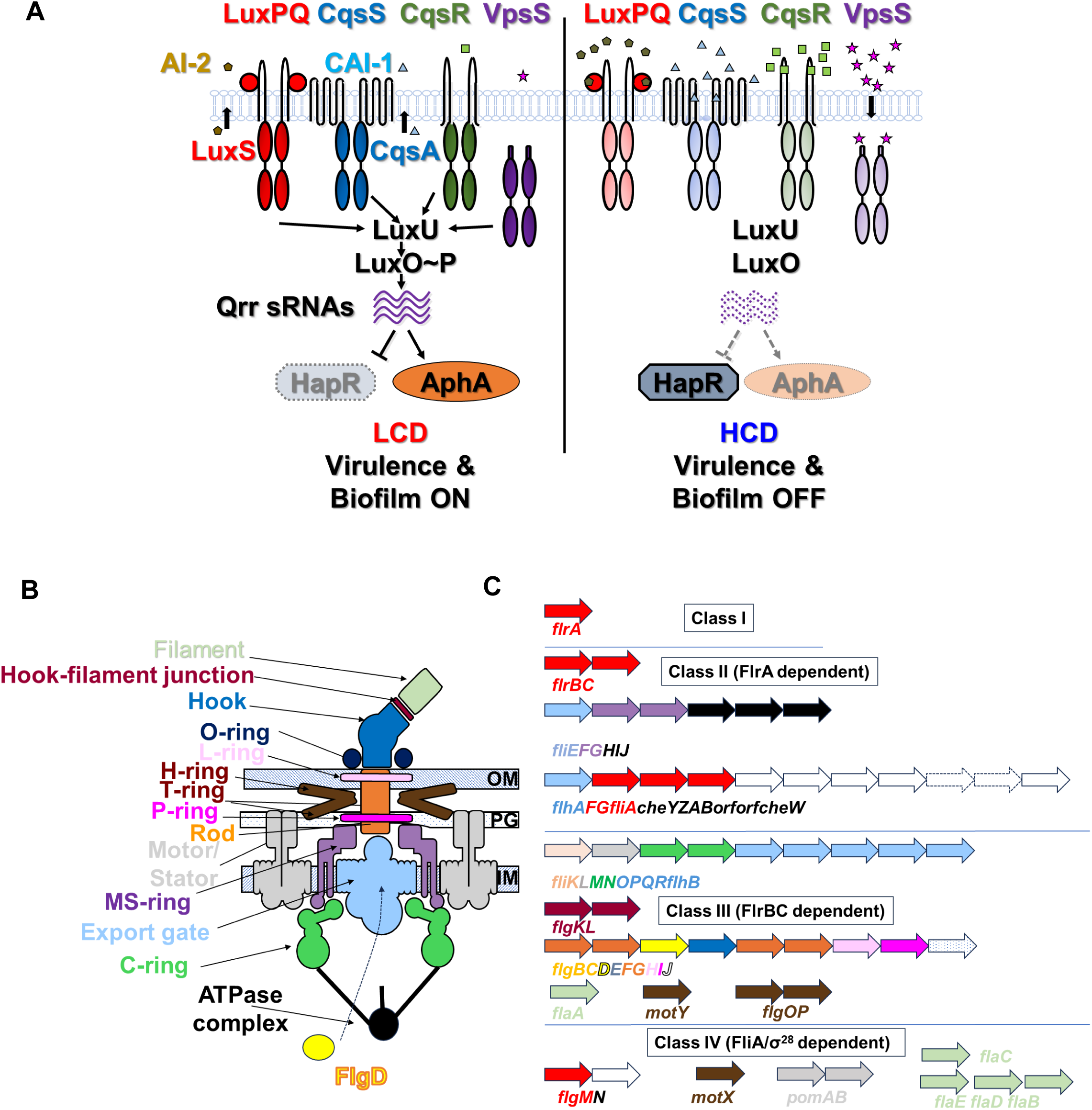
(A) *V. cholerae* LuxO-dependent QS signaling at low (left) and high (right) cell density. LuxO phosphorylation level dictates the amount of the 4 Qrr sRNAs and the overall QS response. (B) Proposed structure of the polar flagellum of *V. cholerae*. The sheath is not shown for clarity and simplicity. Key regulators FlrA, FlrBC, and FliA/FlgM are not shown here with the structure. (C) Hierarchical transcriptional regulation of the four classes of flagellar gene in *V. cholerae*. Red arrows denote genes that encode important flagellar regulators including FlrA, FlrBC, FlgM, and FliA. Arrows with no color are mostly for chemotaxis and not flagellar structural proteins. The rest of the genes are color-coded as the encoded flagellar proteins shown in (B). Genes and proteins are not drawn in scale.

At high cell density (**HCD**), when the extracellular autoinducer concentrations are high, signals binding to the QS receptors inhibits the kinase activity, so LuxO is unphosphorylated. Production of Qrr sRNAs and AphA is halted, and HapR production is activated, resulting in repression of virulence and biofilm genes at HCD. Consistent with this model, mutants lacking LuxO, or all 4 Qrr sRNAs, or all 4 QS receptors, are locked in a HCD QS state and cannot form biofilm [6, 7, 30] and are avirulent [4, 5, 13]. The LuxO-independent DPO/Vqm QS system is also present in *V. cholerae* that regulates virulence and biofilm gene expression [31–34].

While the effects of cell density and autoinducer level on QS signaling are well established, the ways in which additional regulatory systems intersect with the core QS circuitry and how these interactions fine-tune or modulate the overall QS response remain largely unexplored. One such signaling cue may originate from the state of the polar flagellum because the two key QS-regulated processes in the *V. cholerae* life cycle, biofilm formation and host colonization [35–38], are also completely dependent on the functionality of the flagellum. We reason that the polar flagellum is not only essential for motility, chemotaxis, and initial surface attachment, the completeness and integrity of this supramolecular structure could generate additional signals to reshape QS response to coordinate group behaviors under different environments.

The structure and assembly of the polar flagellum in *V. cholerae* (**Fig. 1B**) is similar to the one established in Enterobacteriaceae [35, 36, 39]. Orderly assembly of the flagellum is in part governed by the hierarchical transcription of flagellar genes mediated by several key regulators [35, 36, 39, 40] (**Fig. 1C**). Master regulator FlrA (encoded by the Class I gene *flrA*) governs the expression of a set of Class II flagellar genes which encode the FlrB (histidine kinase) and FlrC (response regulator) to form a two-component system (**TCS**), the MS-ring, several components of the flagellar Type III secretion system (**fT3SS**), the alternative sigma factor (FliA, σ^28^) and some chemotaxis genes [41]. FlrBC in turn activates Class III flagellar genes in *V. cholerae* [42, 43], which encode the remaining subunits of the fT3SS, the P/L rings, rod, hook, the FlaA flagellin, as well as the flagellar hook cap, FlgD. Completion of hook-basal body allows the secretion of anti-σ factor FlgM to relieve the inhibition of σ^28^ which activates the Class IV genes that encode the motor/stator and other flagellins to complete the flagellar assembly [44, 45].

It was previously demonstrated that *hapR* transcription is repressed in *V. cholerae* mutants lacking the flagellar hook cap protein FlgD, and *V. cholerae* cells that have lost their polar flagellum during mucin penetration *in vitro* are locked in LCD QS state [46]. These observations suggest that *V. cholerae* is primed to induce virulence gene expression during initial host colonization through QS repression [46]. However, the exact molecular details of how the flagellum is connected to QS regulation, and the impact of the integrity of the flagellum on other parts of the central QS circuit are both unknown. Since the Qrr sRNAs lie at the heart of the QS circuit (**Fig.1A**) and they directly control many important processes independent of HapR [15, 21, 47–49], we studied if and how the flagellar apparatus regulates *qrr* expression. Through this investigation, we identified that FlgD functions as a previously unknown cue that regulates the activity of FlrBC TCS. Our data suggests that when FlrBC is sufficiently activated in cells with an incomplete flagellum, LuxO is no longer required to produce Qrr sRNAs, thus, QS response becomes independent of cell density. Importantly, the regulatory link between FlgD and QS appears to be evolutionary conserved in multiple *Vibrio* species that uses LuxO as the key QS regulator. Together, our work provides important mechanistic insight on how the input from the flagellum is integrated to control QS-regulated phenotypes outside of its usual role in motility and chemotaxis, bypassing the canonical cell density control.

## RESULTS

### Cell-density independent production of Qrr in mutants lacking flagellar hook cap FlgD

Because transcription of *hapR* is repressed in the Δ*flgD* mutants lacking the flagellar hook cap [46], we tested if defects in flagellar synthesis or regulation also influence *qrr* expression in the Δ*flgD* and other flagellar mutants of *V. cholerae* [35, 36] using a P*_qrr_*_4_-*lux* transcriptional reporter. This bioluminescent reporter has been previously used as a direct and reliable readout of Qrr4 level [13, 14, 50], and thus we could compare the relative light units (RLU, bioluminescence (lux)/OD_600_) produced by these strains over time to monitor their *qrr* expression at different cell densities.

Our analysis revealed that Δ*flgD* mutant lacking the flagellar hook cap was locked in LCD QS state with a mechanism distinct from the one previously described [46]. At HCD (OD_600_ ∼1), a condition where P*_qrr_*_4_-*lux* expression was usually low in wildtype (**WT**) *V. cholerae* C6706, P*_qrr_*_4_-*lux* expression was exceptionally high in the Δ*flgD* mutant (**Fig. 2A-B**). The effect of the Δ*flgD* mutation on P*_qrr_*_4_-*lux* expression could be complemented by another copy of *flgD* under control of the native P*_flgB_* promoter (**Fig. 1C, 2A-B**). With additional P*_qrr_*-*lux* reporters, we found that Δ*flgD* mutant produced high level of all four Qrr sRNAs at HCD (**Supplementary Fig. 1**). The same phenotype was observed in the Δ*flgD* mutant of another *V. cholerae* strain E7946 (**Fig. 2B**), suggesting *qrr* repression by FlgD is a universal regulatory mechanism in multiple *V. cholerae* strains.

**Fig. 2.**
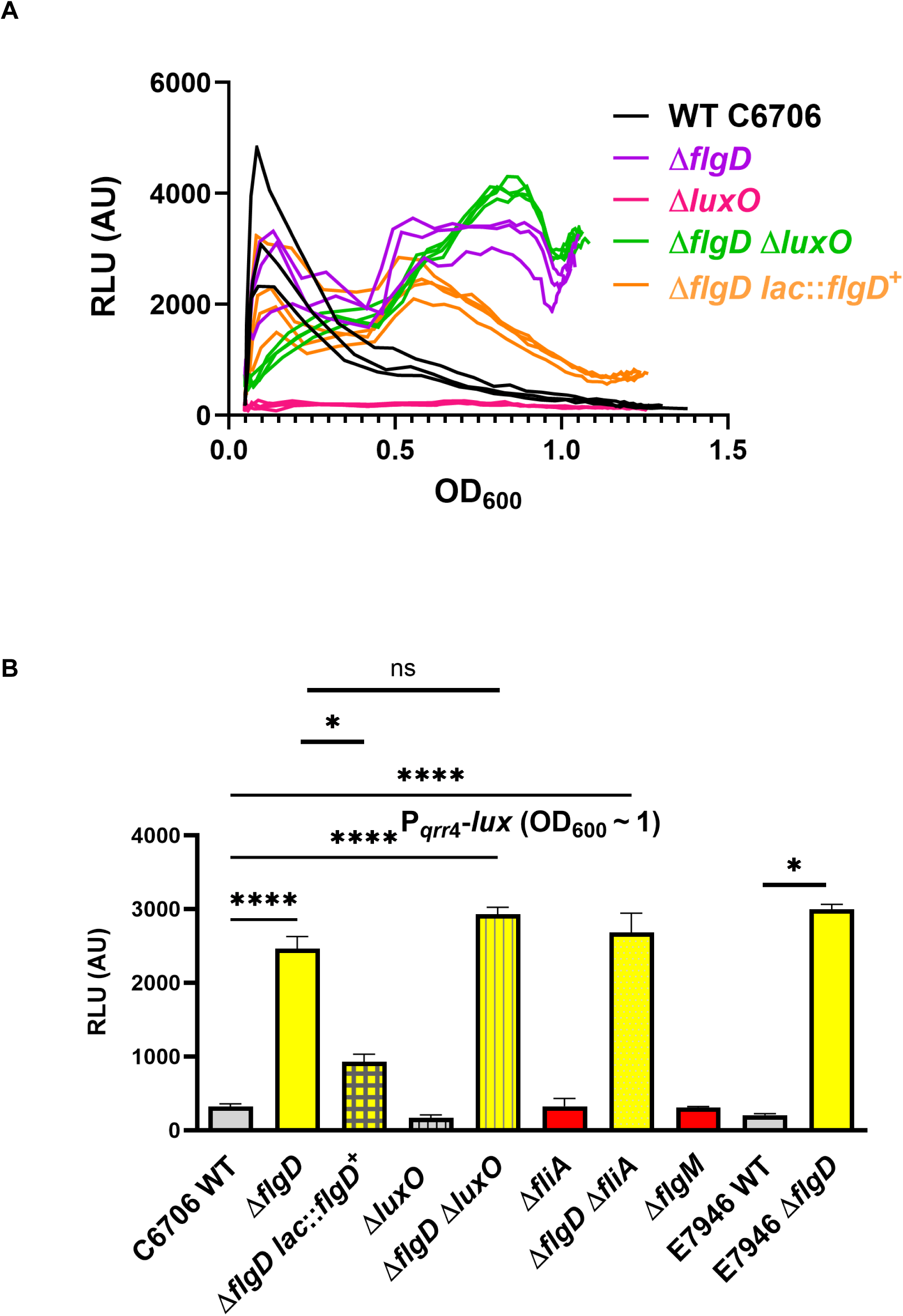
*qrr*4 transcription measured by a bioluminescence reporter (P*_qrr_*_4_-*luxCDABE*) in different strains during growth (**A**) and at HCD (OD∼1) (**B**). Representative results with multiple replicates (n >3) are shown. Three technical replicates are shown to show variation in (A) as OD_600_ at each time point cannot be directly averaged in different samples. Ordinary one-way ANOVA with Dunnett’s multiple-comparison test was used for comparison (*denotes p<0.05; ns, p>0.5)

It was previously shown that a high level of anti-σ factor FlgM is aberrantly secreted by the Δ*flgD* mutant, increasing the intracellular activity of FliA (σ^28^), resulting in *hapR* transcriptional repression with an unknown mechanism [46, 51]. However, we found that the effect of the Δ*flgD* mutation on *qrr* transcriptional activation was independent of FliA because the Δ*flgD* Δ*fliA* double mutant still produced high level of P*_qrr_*_4_-*lux* and the Δ*flgM* mutant still produced low level of P*_qrr_*_4_-*lux* at HCD (**Fig. 2B**), indicating the effect of Δ*flgD* mutation on *hapR* transcriptional repression and *qrr* activation is through two distinct mechanisms. Thus, the Δ*flgD* mutation could decrease both *hapR* mRNA and HapR protein amounts (by increasing Qrr, **Fig.1A**), enforcing a locked LCD QS state.

## Mutants lacking FlgD produce Qrr and regulate multiple QS behaviors independent of LuxO

All *V. cholerae* mutants lacking LuxO assayed to date exhibit phenotypes identical to those lacking all four Qrr sRNAs [13, 22, 47] (**Fig. 1A**). Thus, the current prevailing view is that phosphorylated LuxO (LuxO∼P) is absolutely indispensable for *qrr* expression. Consistent with this view, P*_qrr_*_4_-*lux* expression was undetectable in the Δ*luxO* mutant at all cell densities (**Fig. 2A-B**), surprisingly, the Δ*flgD* Δ*luxO* produced high P*_qrr_*_4_-*lux* activity, suggesting increased production of Qrr in the Δ*flgD* mutant at HCD is completely independent of LuxO.

To test if QS-regulated gene expression is also uncoupled from LuxO regulation in the Δ*flgD* mutant, we measured the expression of the VPS-II operon which is repressed by HapR (i.e., activated by Qrr) [52] using the P*_vpsL_*-*lux* reporter. Again, we found that P*_vpsL_*-*lux* was undetectable in the Δ*luxO* mutant, suggesting VPS-II gene expression was highly repressed in the Δ*luxO* mutant that constantly makes HapR (**Supplementary Fig. 2**). However, P*_vpsL_*-*lux* expression was restored in the Δ*luxO* Δ*flgD* mutant (**Supplementary Fig. 2**), suggesting Δ*flgD* not only affects *qrr* expression, but this flagellar mutation also imposes a downstream regulatory effect on the expression of other QS-controlled genes.

We then tested if Δ*flgD* mutation affects other LuxO/Qrr-controlled phenotypes. We previously discovered that mutants constantly producing Qrr are unable to alter the pyruvate flux to convert fermentable carbon sources into neutral acetoin and 2,3-butanediol molecules to offset organic acid production. As a consequence, these strains rapidly lose viability when grown with fermentable carbon sources [21]. Consistent with our previous findings, WT and Δ*luxO* were viable and grew comparably during glucose fermentation, in contrast, the Δ*flgD* and Δ*flgD* Δ*luxO* mutants, producing high levels of Qrr, lost their viability under the same growth conditions (**Fig. 3A**). There was no difference in viability among these stains when they were grown with non-fermentable glycerol (**Fig. 3A**).

**Fig. 3.**
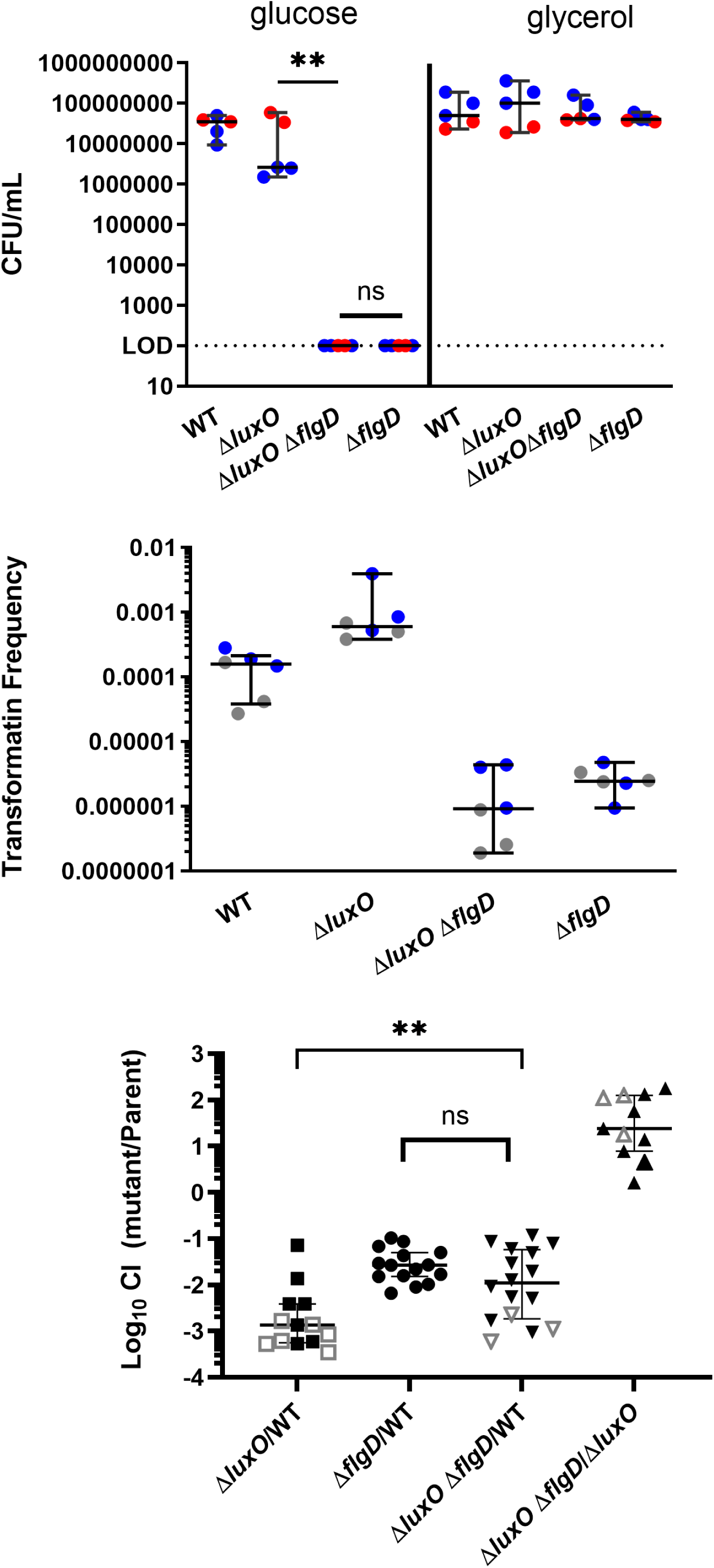
Δ*flgD* mutation bypasses LuxO regulation of several QS behaviors. (**A**) Δ*flgD* mutation is epistatic to Δ*luxO* in determining cell viability during glucose fermentation. LOD is the limit of detection for viable cell count (**B**) Δ*flgD* mutation is epistatic to Δ*luxO* in affecting natural competence development induced by chitin. Transformation frequency is calculated by dividing the number of transformants to the number of total viable cells and the amount of DNA. In A and B, results from two biological replicates are shown in different colors. **(C)** Infant mouse small intestinal colonization competition of different *V. cholerae* strains (∼8 mice were tested in each competition and two competitions were performed). Each symbol represents the competitive index (CI) in an individual mouse, and horizontal lines indicate the median with interquartile range. Open symbols represent data below the limit of detection for the Δ*luxO* mutants. In such case, it was assumed that there was 1 CFU present at the next lowest dilution to calculate the CIs. Mann-Whitney test was used for comparison (*denotes p<0.05; ns, p>0.5).

Another well-studied QS-regulated phenotype is the development of genetic competence, which is activated by HapR (i.e., repressed by Qrr) in the presence of chitin [12, 53]. To test how Δ*flgD* mutation affects the development of genetic competence, we measured the efficiency of natural transformation of different Δ*luxO* and Δ*flgD* mutants. As expected, as both WT and the Δ*luxO* mutant are capable of producing HapR, they displayed a similar efficiency in transformation (**Fig. 3B**). In contrast, the Δ*flgD* and Δ*flgD* Δ*luxO* mutants, producing high Qrr and thus low HapR, were >100-fold less efficient in transformation than WT and the Δ*luxO* mutant (**Fig. 3B**). Therefore, the Δ*flgD* mutation is epistatic and it reverses the original glucose fermentative sensitivity and genetic competence phenotypes displayed by the Δ*luxO* mutants.

We further investigated the *in vivo* role of the new connection between FlgD and QS using an infant mouse intestinal colonization model [54]. As shown before and here, due to the inability to produce the key colonization factor TCP [4, 13], the *ΔluxO* mutants were highly defective in host colonization (∼3-log decrease, **Fig. 3C**) and sometimes they were completely outcompeted by WT and were undetectable *in vivo* (open symbols in **Fig. 3C**). In contrast, host colonization by the Δ*flgD* mutant was only attenuated and outcompeted by 1- to 2-log [46], and Δ*flgD* mutants could be recovered from the infected intestines (**Fig. 3C**). We hypothesized that the Δ*flgD* mutation could restore *qrr* expression and TCP production and thus compensate for the colonization defects of the Δ*luxO* mutants. Indeed, similar to the Δ*flgD* mutant, the Δ*flgD* Δ*luxO* double mutants were only outcompeted *in vivo* by WT by 1- to 2-log (**Fig. 3C**). Moreover, when competed against the Δ*luxO* mutants, the Δ*flgD* Δ*luxO* double mutants colonized 1- to 2-log better (**Fig. 3C**). Our results suggest a signal from an incomplete flagellum is integrated to bypass LuxO to activate virulence gene *in vivo*.

The results of these *in vitro* and *in vivo* phenotypic characterizations suggest that the connection between the flagellum and QS has broad impact on *V. cholerae* group behaviors: Δ*flgD* mutation not only affects Qrr levels, but this specific flagellar defect overrides the dependency of the canonical LuxO control on QS-regulated gene expression and impacts the physiology of *V. cholerae*. To our knowledge, this is the first instance where *qrr* expression and QS response is totally uncoupled from the dependence of LuxO, and thus QS response can become independent of cell-density control when the flagellum assembly is disrupted.

## FlgD represses *qrr* expression in multiple *Vibrio* species

Many *Vibrio* species use LuxO to activate *qrr* expression to control QS [55–60]. To test if Δ*flgD* mutation also eliminates the requirement of LuxO to activate *qrr* transcription in these *Vibrio* species, we introduced the Δ*flgD* mutation in *V. campbellii* (previously classified as *V. harveyi* [61])*, V. parahaemolyticus*, and *V. vulnificus*, and measured *qrr* expression using the same P*_qrr4_-lux* reporter either with and without the Δ*luxO* mutation, or with and without a compound (**530A**) that has been previously shown to inhibit LuxO activity in *V. cholerae* and *V. parahaemolyticus* [62, 63]. Similar to *V. cholerae,* in the WT strain backgrounds, P*_qrr_*_4_-*lux* activity was high at LCD in these *Vibrio* species and low at HCD (**Fig. 4**), suggesting LuxO is activated at LCD. As expected, in *V. campbellii*, the Δ*luxO* mutation significantly decreased P*_qrr_*_4_-*lux* expression at all cell densities. The Δ*flgD* mutant displayed slightly higher P*_qrr_*_4_-*lux* activity at HCD compared to WT. But importantly, the Δ*flgD* Δ*luxO* mutant displayed similar P*_qrr_*_4_-*lux* activity as the Δ*flgD* mutant, suggesting LuxO was not required for the *V. campbellii* Δ*flgD* mutant to express *qrr* (**Fig. 4A**). In *V. parahaemolyticus* and *V. vulnificus,* Δ*flgD* mutant likewise produced more Qrr at HCD compared to WT (**Fig. 4B-C**). While the LuxO inhibitor 530A abolished *qrr* expression in the WT strain of these two *Vibrio* species, the Δ*flgD* mutants of these two species were more resistant to the inhibition and P*_qrr_*_4_-*lux* expression remained high (**Fig. 4B-C**). These results suggest that FlgD is involved in repression of *qrr* expression and LuxO is not required for *qrr* expression in multiple *Vibrio* mutants lacking FlgD, and thus the regulatory connection between flagellum, in particular the Δ*flgD* mutation, and QS is evolutionarily conserved among many *Vibrio* species.

**Fig. 4.**
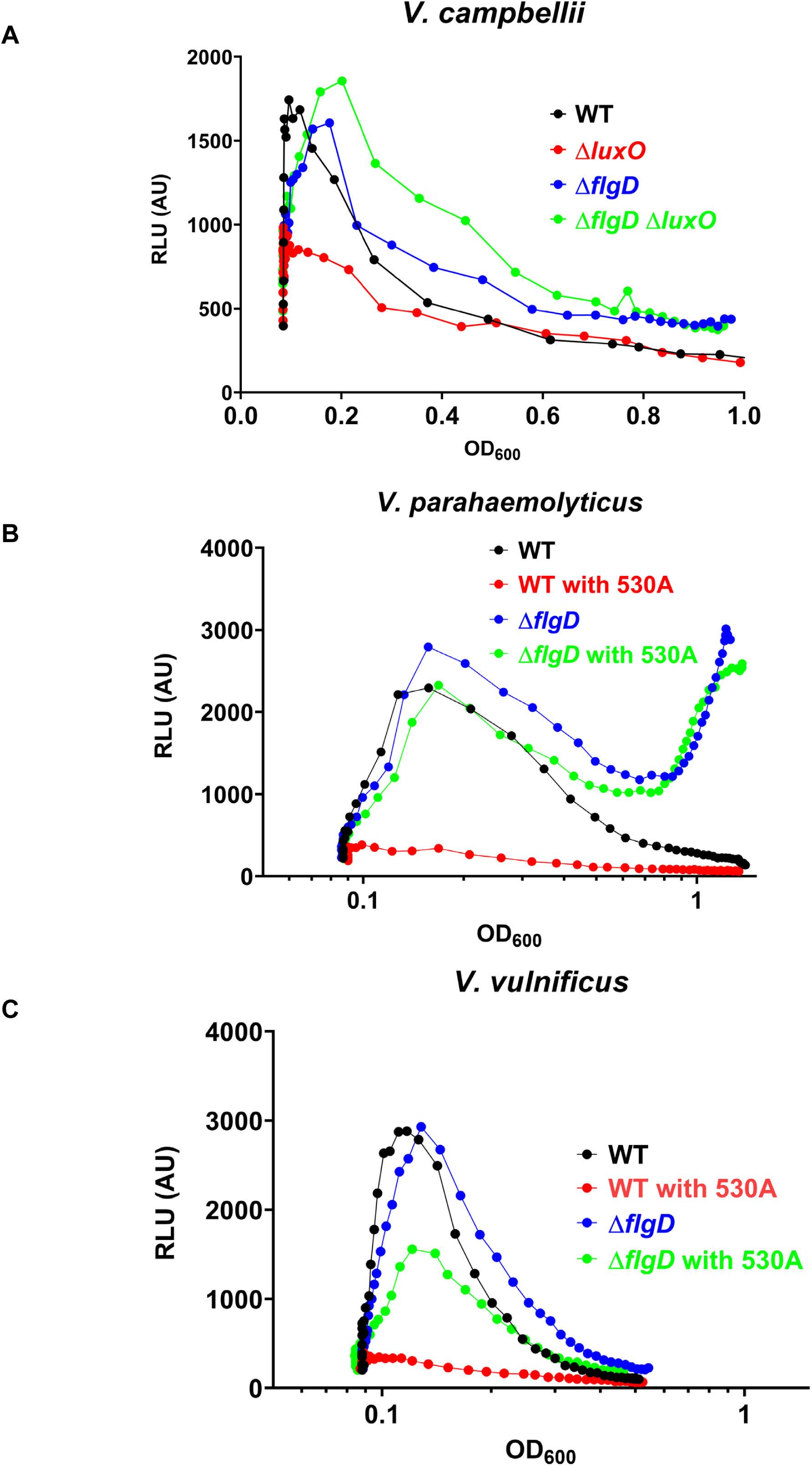
Transcriptional regulation of *qrr* in different *Vibrio* species. **(A)** *qrr* transcription pattern in *V. campbellii* strains with and without FlgD and/or LuxO. **(B and C)** *qrr* transcription patterns in *V. parahaemolyticus* (B) *and V. vulnificus* (C) strains with and without FlgD and/or LuxO chemical inhibitor 530A. Experiments were repeated at least 3 times, representative results are shown.

### The FlrBC two-component system is required for high *qrr* expression in the Δ*flgD* mutant

To understand the molecular basis by which the Δ*flgD* mutation increases *qrr* expression, we measured *qrr* expression in additional *V. cholerae* flagellar mutants using the P*_qrr4_*-*lux* reporter. To generate this panel of mutants, we used two approaches. First, we introduced deletion mutations by using allelic exchange [64] to generate strains carrying in-frame deletions (Δ) in different flagellar genes (**Fig. 5A**). Second, we used PCR-amplified transposon insertion mutations in various flagellar genes from an ordered transposon mutant library [65] and introduced them into our assay strains by natural transformation [66]. The transposon insertions in these strains were subsequently removed by inducing the production of FLP recombinase encoded on a plasmid, leaving an 192-bp in-frame FRT insertion (::*FRT*) within the gene [65, 67]. This FLP plasmid was then cured before the bioluminescence assays (**Fig. 5B**). For the several strains we have characterized, we did not observe any difference in P*_qrr4_*-*lux* activity regardless of which method was used to generate the mutation in the same gene.

**Fig. 5.**
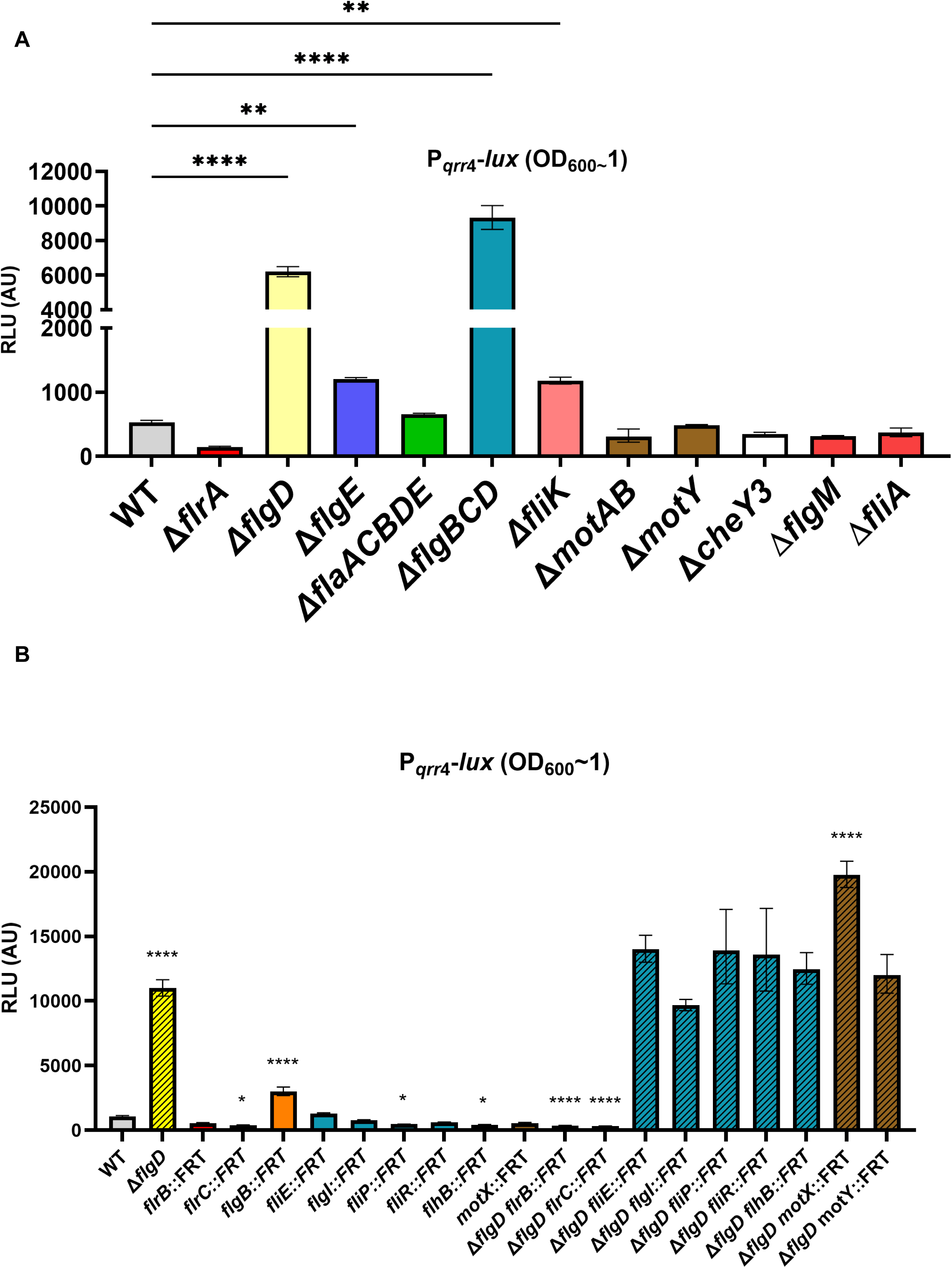
*qrr* expression in different *V. cholerae* flagellar mutants. P*_qrr_*_4_-*lux* expression patterns in different *V. cholerae* flagellar mutants at OD_600_∼1. **(A)** The top panel shows the strains constructed using allelic exchange with in-frame deletions. **(B)** The bottom panel shows the strains constructed with in-frame FRT insertions in WT or in the Δ*flgD* background. Expression was measured in these strains at the same time if they are shown in the same panel. Experiments were repeated at least 3 times, representative results are shown. For statistical analysis, expression patterns of mutants constructed in WT background were compared with that from WT, and expression patterns of mutants constructed in the Δ*flgD* background were compared with that from the Δ*flgD* mutant.

When these flagellar mutations were introduced to the WT C6706, most of the mutations did not significantly increase P*_qrr_*_4_*-lux* expression at HCD as observed in the Δ*flgD* mutant. These include mutations that eliminate some or all the flagellins (Δ*flaACBDE*), the flagellar hook (Δ*flgE*), the motor/stator, rings (Δ*motXY*, Δ*motAB* (*pomAB), flgI:: FRT*), chemotaxis (Δ*cheY3*), the fT3SS portal (*fliR::FRT*, *fliP::FRT*, *flhB::FRT*), proximal rod protein (*fliE::FRT)*, the “molecular ruler” (Δ*fliK*), and all the key flagellar regulators (Δ*flrA*, *flrB::FRT, flrC::FRT,* Δ*fliA,* Δ*flgM*) (**Figs. 5A-B**). As mutants lacking some of these flagellar components (e.g., FlaA, MotXY) have been previously found to have altered cyclic di-GMP levels [37], the lack of *qrr* induction in these mutants suggest that the regulatory link between Δ*flgD* and QS is not directly related to the change of the c-di-GMP levels induced by flagellar defects. We also noticed the Δ*flgBCD* and *flgB::FRT* mutants also displayed higher P*_qrr_*_4_*-lux* activity at HCD (**Figs. 5A-B**), however, ectopic expression of *flgD* in the Δ*flgBCD* mutant repressed P*_qrr_*_4_*-lux* activity (**Supplementary Fig. 3**), suggesting the Δ*flgB* mutation on *qrr* expression is indirect; and the mutation likely aberrantly affects FlgD level inside the cell. Together, our results suggest FlgD functions as a specific signal that connects the polar flagellum to QS via *qrr* regulation.

Given FlgD is needed for repression of *qrr* expression at HCD, the low *qrr* expression displayed by the Δ*flrA, flrB*::*FRT,* or *flrC*::*FRT* mutants was unexpected because these mutants should also express low level of *flgD,* which is a Class III gene [41, 45] (**Fig. 1B**). To understand the underlying mechanisms, we determined how these flagellar mutations affect *qrr* expression in combination with the Δ*flgD* mutation. Because *flrA*, a Class I gene, is expected to be expressed in the Δ*flrBC* mutant, we focused only on *flrB* and *flrC* mutations. We found that either *flrB*::*FRT* or *flrC*::*FRT* mutations introduced in the Δ*flgD* mutant displayed low P*_qrr_*_4_-*lux* activity at HCD (**Fig. 5B**), indicating that absence of either *flrB* or *flrC can* suppress the high *qrr* expression in Δ*flgD*.

As shown above, Δ*fliA* mutation (σ^28^ deficient) did not suppress high *qrr* expresion in the Δ*flgD* mutants (**Fig. 2B**), indicating none of the components encoded by the Class IV genes (e.g., FlgMN, motor/stator, flagellins, **Fig.2C**) is involved. Therefore, we focused on the components encoded by the FlrBC-dependent Class III genes. None of the Class III mutations we tested (*fliR::FRT*, *fliP::FRT*, *flhB::FRT*, *flgI::FRT*, *motY::FRT*) could suppress the increased *qrr* expression in the Δ*flgD* mutant (**Fig. 5B**), therefore, we reasoned that high *qrr* expression in the Δ*flgD* mutant is dependent on the FlrBC itself, rather than the genes controled by this TCS.

### FlrC directly activates *qrr* transcription by utilizing LuxO binding sites in the Δ*flgD* mutant

FlrB is a cytoplasmic histidine kinase carrying a PAS sensing domain and localized to the pole [43, 68, 69]. It phosphorylates FlrC to activate this σ^54^-dependent response regulator for Class III gene expression. We hypothesized that the FlrBC TCS could be activated in the Δ*flgD* mutant and that activated FlrC drives *qrr* expression. To test this hypothesis, first, we produced the consitutively active FlrC^M114I^ variant [42, 43] in the *flrB*::FRT*, flrC*::FRT, or Δ*flgD flrC*::FRT mutants of *V. cholerae,* and we found that FlrC^M114I^ alone was sufficient to induce P*_qrr_*_4_*-lux* expression in these strains (**Fig. 6A**). Moreover, production of FlrC^M114I^ alone in the heterologous host *E. coli,* which does not encode this TCS, was also sufficient to drive P*_qrr_*_4_*-lux* expression (**Fig. 6B**). Indeed, when the FlrBC of *V. cholerae* was introduced to *E. coli* MG1655, P*_qrr_*_4_*-lux* expression was highly induced compared to the empty vector control, suggesting FlrBC directly regulates *qrr* expression in the absence of other *V. cholerae* QS components (**Fig. 6C**). Together, our data suggest FlrBC, when activated under specific circumstances, can also modulate QS response.

**Fig. 6.**
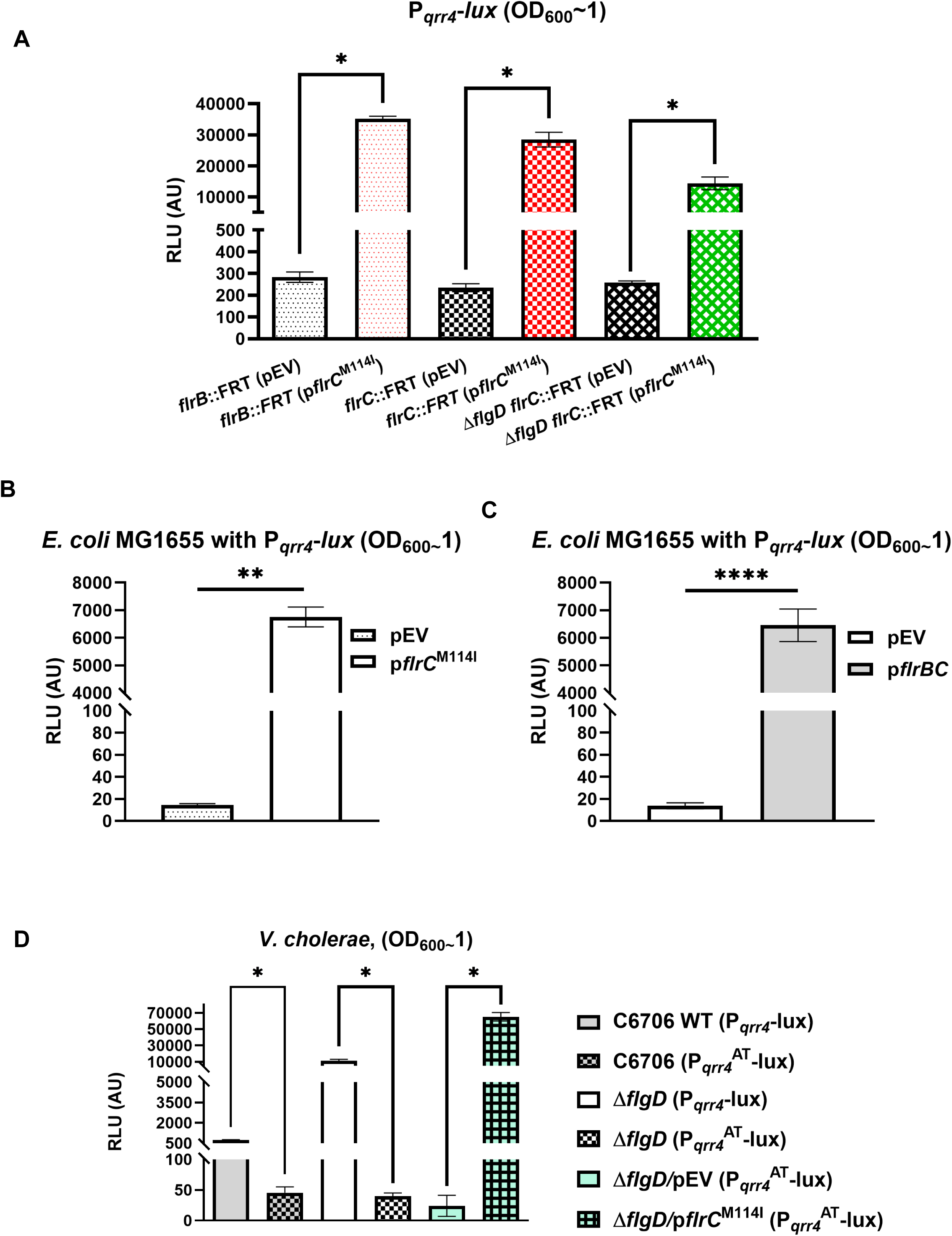
FlrC directly activate *qrr* transcription. **(A)** Overexpression of a hyperactive constitutive mutant of FlrC (*flrC*^M114I^) under an inducible promoter activates *qrr* transcription in *V. cholerae* (C6706) flagellar mutants lacking *flrB* and *flrC* in both WT and in Δ*flgD* mutants. **(B)** Overexpression of a hyperactive constitutive mutant of FlrC (*flrC*^M114I^) under an inducible promoter activates *qrr* transcription in the heterologous *E. coli* host (MG16555). **(C)** Expression of *V. cholerae flrBC* under an inducible promoter promotes *qrr*4-*lux* expression in *E. coli* strain MG1655. **(D)** Mutation in the known LuxO binding-site on the *qrr*4 promoter (P ^AT^) abolishes both LuxO- and FlrC-mediated *qrr-lux* expression in *V. cholerae* WT and Δ*flgD* mutants, whereas hyperactive mutant of *flrC*^M114I^ drives *qrr* transcription from the mutated P ^AT^ promoter. Statistical significance was determined using Mann-Whitney (6A, 6D) test and ordinary two-way ANOVA (6B, 6C) (*denotes p<0.05).

We also tested the requirement for FlrC binding on the P*_qrr4_* promoter. FlrC and LuxO are both members of the NtrC family of σ^54^-dependent response regulators, raising the possibility that FlrC may activate *qrr* transcription by binding to the recognition sites for LuxO within the *qrr* promoter. Based on previously defined LuxO-binding sites in the *qrr* promoters [22], we mutated one of the two known LuxO-binding sites in our reporter construct (named P*_qrr4_*^AT^) **(Supplementary Fig. 4)**. We found that mutating this LuxO-binding site strongly reduced P ^AT^*-lux* expression in both WT and the Δ*flgD* mutants (**Fig. 6D**). This promoter is not completely defective as hyperactive FlrC^M114I^ was active in promoting *qrr*4 expresison with this mutated promoter, suggesting RNA polymerase/σ^54^ can bind to this mutated promoter (**Fig. 6D**). These new results suggest that activated FlrC in the Δ*flgD* mutant is a key positive regulator of *qrr* expression and support the idea that activated FlrC directly transcribes Qrr sRNAs.

## Cytoplasmic FlgD functions as an inhibitor of FlrBC two-component system

Because *qrr* expression was elevated in the absence of FlgD, and this activation occurred through the FlrBC TCS, suggesting the possibility that FlgD inhibits FlrBC signaling (**Fig. 7A**). The exact nature of the input signal for FlrBC is unkown so it is unclear how FlrB activity is regulated [69]. To drive Class III flagellar gene expression, full activity of FlrBC requires many components of the *V. cholerae* fT3SS (Export gate, MS Ring, C ring, and ATPase) and, thus, the fT3SS is proposed to function as a regulatory checkpoint during flagellar biosynthesis [70]. However, we found that mutations that abolish the function of fT3SS (*flhB::FRT*, *fliP::FRT*, *fliR::FRT*) could not suppress the high *qrr* expression in the Δ*flgD* mutant (**Fig. 5B**), suggesting fT3SS does not function as a direct positive regulator of FlrBC on *qrr* regulation. Instead, since fT3SS mutants is defective in flagellar secretion [40, 71], presumably causing the accumulation of many flagellar proteins including FlgD in the cytoplasm (**Fig. 7A**), we hypothesized that fT3SS functions as a regulatory checkpoint by secreting a cytoplasmic inhibitor of FlrBC, and based on our results the most likely candidate for this inhibitor is FlgD (**Fig. 7A**).

**Fig. 7.**
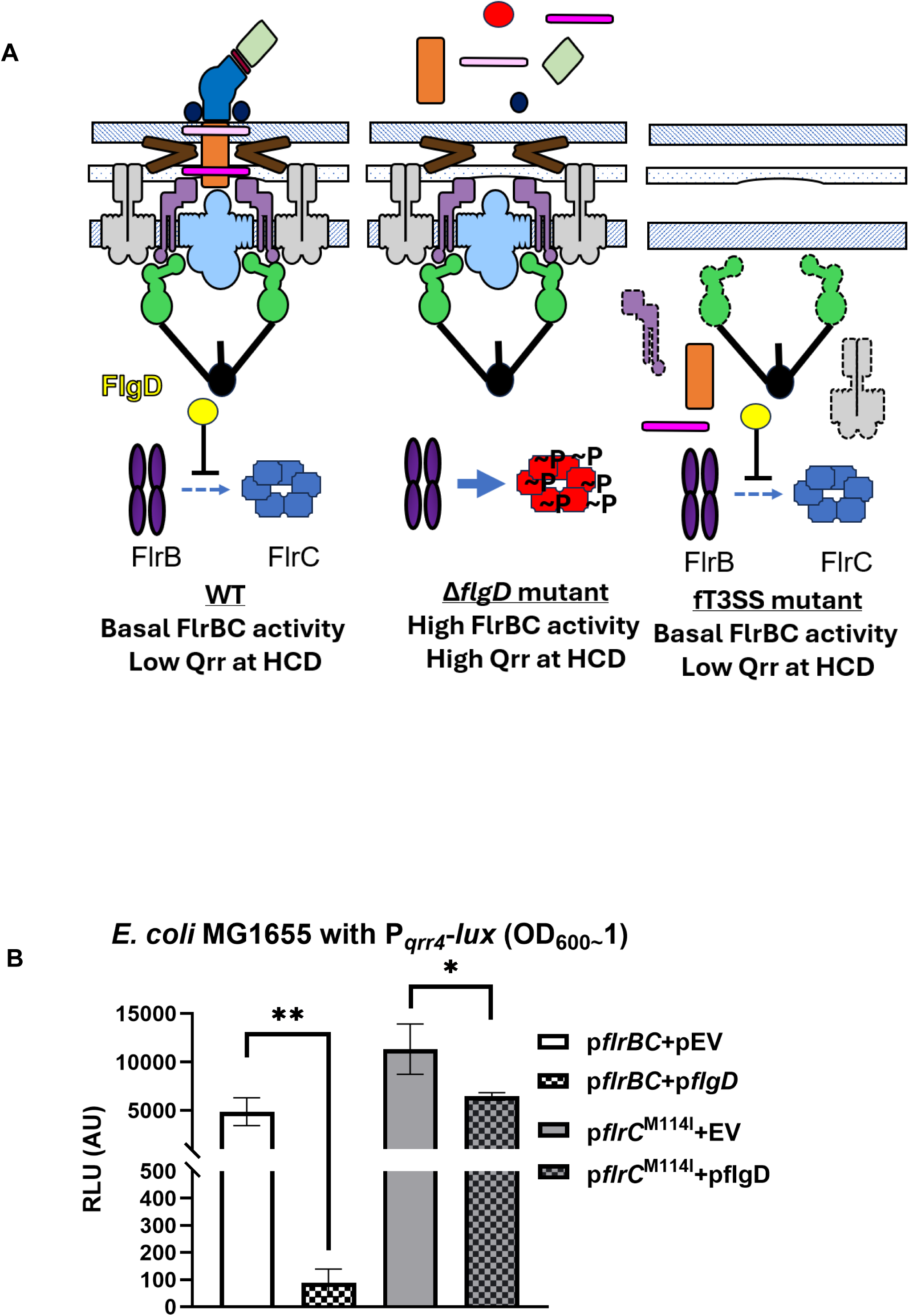
Cytoplasmic FlgD is the inhibitor of FlrBC to regulate *qrr* expression. (A) Proposed model of how flagellar defects in various mutants affect FlrBC activity via FlgD. Structures are color coded as in Fig. 2. Sheath is not shown. Δ*flgD* mutants release many flagellar proteins outside of the cell. Many flagellar components are not secreted nor assembled in the fT3SS mutant, leading to accumulation in the cytoplasm. (B) Effect of FlgD expression on P*_qrr_*_4_-*lux* expression in *E. coli* strain expressing *V. cholerae* FlrBC and FlrC^M114I^. Mann-Whitney test was used for comparison (*denotes p<0.05).

The idea that FlgD, but not other flagellar proteins, as an inhibitor of FlrBC is supported by the observations where FlrBC-dependent P*_qrr_*_4_-*lux* expression in the heterologous *E. coli* host was strongly repressed (∼50-fold) by the *V. cholerae* FlgD (**Fig.7B**), but *V. cholerae* FlgD imposed almost no inhibition (< 2-fold) of P*_qrr_*_4_-*lux* expression in the *E. coli* strain expressing the constitutively active FlrC^M114I^ (**Fig. 7B**).

To further understand the regulatory role of FlgD on controlling FlrBC activity, we used a bacterial two-hybrid system (BACTH) [72] to identify potential protein-protein interactions among FlgD, FlrB, and FlrC inside bacterial cells. We constructed protein fusions of FlgD, FlrB, and FlrC to the complementary fragments of the *Bordetella pertussis* adenylate cyclase, known as T25 and T18, in the adenylate cyclase deficient *E. coli* strain BTH101. If any of the two proteins interact within the cell, functional adenylate cyclase will form, leading to cAMP synthesis and activation of the reporter *lacZ* gene, which can be scored as blue colonies on LB plates with an X-Gal indicator. As expected, since FlrB, FlrC and FlgD all form multimers on their own [68, 73–75], blue colonies were obtained when T25-FlrB and T18-FlrB (or T25-FlrB and FlrB-T18), T25-FlrC and T18-FlrC, or T25-FlgD and FlgD-T18 were co-produced in BTH101, in contrast, no blue colonies were identified with the empty vector negative controls (**Fig. 8A**).

**Fig. 8.**
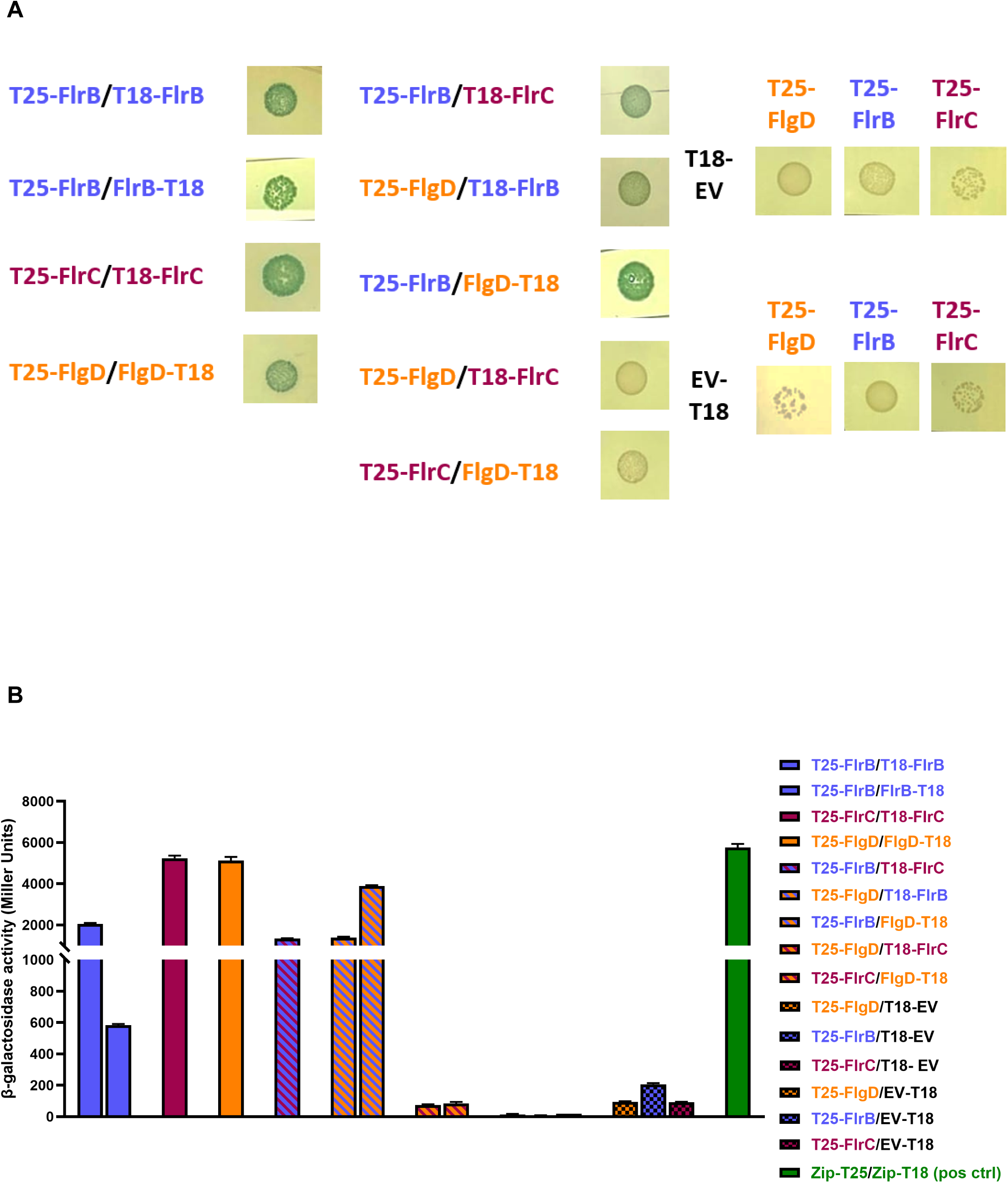
BATCH analysis of protein-protein interactions among FlgD, FlrB, and FlrC. (A) Colony morphology of strains expressing FlgD, FlrB, and FlrC fused to different T25 and T18 combinations. Morphologies were imaged on LB agar plates supplemented with X-Gal and IPTG. EV denotes empty vector. **(B)** Quantitative β-galactosidase assay corresponding to the BACTH interactions shown in panel A. pos ctrl denotes positive control.

Using these functional protein fusions, we then tested the interactions among each other. Consistent with the predicted signaling functions, we detected blue colonies when T25-FlrB and T18-FlrC were co-produced in BTH101 (**Fig. 8A**). Finally, blue colonies were identified when T25-FlgD and T18-FlrB or when T25-FlrB and FlgD-T18 were co-produced, suggesting FlgD interacts directly with FlrB. In contrast, we did not detect any blue colonies when T25-FlgD and T18-FlrC or T25-FlrC and FlgD-T18 were co-produced (**Fig. 8A**), suggesting FlgD does not directly interact with FlrC. These interactions were further quantified using β-galactosidase assays, and the results corroborated that FlgD and FlrB interact within bacterial cells (**Fig. 8B**). These results suggest that FlgD directly interacts with FlrB that could lead to inhibition of FlrBC signaling and prevents FlrC activation (**Fig. 7A**).

As *flgD* is one of the Class III flagellar genes regulated by FlrBC (**Fig.1B**), we propose the level of FlgD inside the cell is sensed by FlrB through direct protein-protein interaction; and through this feedback mechanism Class III flagellar genes are regulated according to the completeness of the flagellum. Our model also illustrates a novel LuxO-independent activation mechanism on *qrr* expression caused by a specific defect in the flagellum that depends on the FlrBC TCS by sensing the levels of FlgD in the cytoplasm.

## Discussion

For decades, the established function of QS is for cell-density/autoinducer sensing [1, 2]. In contrast, our knowledge of non-chemical factors that influence QS is very limited. In a seminal study, the Zhu Lab demonstrated that *hapR* transcription is repressed in *V. cholerae* cells that have lost their flagellum during mucin penetration *in vitro*, and they postulated that *hapR* repression primes the pathogen to enter a LCD QS state and induce virulence gene expression during initial host colonization [46]. However, the mechanism underlying the connection between QS and flagellum, and how the dynamic of QS response is tied to the flagellar assembly/disassembly has been elusive.

Here we present evidence that disruption of the flagellar apparatus bypasses the requirement for the canonical regulator, LuxO, in driving *qrr* expression in *V. cholerae* and related *Vibrio* species using a mechanism distinct from the previous study [46]. Our results indicate that FlrBC activity is critical for this connection and we identifed that the amount of cytoplasmic FlgD functions as a key regulator that directly modulates FlrBC activity. Our study provides an explanation for the previously unclear mechanism governing FlrBC signaling [69, 70].

Flagellar proteins are secreted by the fT3SS in order and FlgD is an early fT3SS substrate [39]. Once the hook-basal body assembly is completed, a switch in specificity occurs for secretion of late-type substrates to complete the flagellar biosynthesis [39]. Therefore, we hypothesize that cytoplasmic FlgD acts as an indicator of the completeness of the flagellar structure (**Fig. 7A**). In cells with an intact flagellum, FlgD secretion should have already completed, and FlrBC activity is kept at a basal level by the cytoplasmic FlgD that has been made. This feedback mechanism would also ensure FlrBC-dependent Class III genes (encoding rod, hook, P/L-rings, etc.) are not unnecessarily expressed. In contrast, in cells with an incomplete flagellar strcuture, cytoplasmic FlgD could be secreted via fT3SS that leads to activation of FlrBC (**Fig. 7A**), allowing restoration of Class III gene reactivation. This proposed model is consistent with previous studies on flagellar gene regulation [35, 36, 41–43, 69, 70, 76]. Besides FlrBC, control of flagellar gene expression is commonly mediated by TCS signaling in Gram-negative bacteria with a polar flagellum (e.g., FleSR in *Pseudomonas* and FlgSR in *Campylobacter jejuni*) [70, 77–80]. For some of these flagellar signaling pathways, the presence of the fT3SS appears to function as the key regulatory checkpoint [70, 78]. It remains to be determined if the role of fT3SS in the regulation of these TCSs is due to the secertion of FlgD or other flagellar proteins that act as the feedback inhibitory signals.

Outside of the role of FlrBC in flagellar transcriptional hierarchy, we also reveal an additional layer of regulation that suggests the structural components of the flagellum can actively modulate QS pathways independent of the canonical cell density control to promote LuxO-independent *qrr* activation and regulate QS-controlled phenotypes (**Figs. 2-7**). We reasoned that *V. cholerae* (or other *Vibiro*) will most likely lose the flagellum when moving in viscous environments (e.g. after mucin penetration [46]) or approaching a solid substratum, not only the loss of flagellum could trigger an increase in c-di-GMP production to promote biofilm formation [37], our findings suggest that flagellum loss leads to FlrC-dependent/LuxO-independent activation of *qrr* to license these cells to enter into a sessile/virulent LCD QS mode, increasing bioiflm/virulence gene expression to increase the overall fitness.

Several key questions arise from this study: It is still unclear how FlgD regulates FlrB activity. The hook cap protein could directly interact with FlrB and inhibit its autophosphorylation, and/or its phosphotransfer to FlrC. Moreover, FlrB is localized to the pole [69]. Therefore, it is possible the absence and presence of FlgD affects FlrB localization and its activity. Both LuxO and FlrC belong to the family of bacterial enhancer binding proteins (**bEBPs**) that works with σ^54^-holoenzyme for gene regulation [81]. Like most bEBPs, LuxO binds ∼100 bp upstream of the σ^54^ promoter of *qrr*1-4 [82]. In contrast, FlrC binds >100 bp downstream of the σ^54^ promoter (an enhancer site) to promote *flaA* transcription [42]. However, our transcriptional reporter for measuring *qrr* expression contains only the σ^54^ promoter and the known upstream LuxO-binding sites of *qrr*4. Thus, activated FlrC likely uses a mechanism to activate *qrr* different from how it activates *flaA*. Further experimentation is requried to understand these machanisitc details.

What is the physiological role for this new regulatory connection? It was shown that *Vibrio* spp. eject their flagella at the base of the hook when nutrients are limited [83, 84]. The dynamic nature of flagellar assembly/disassembly in *Vibrio* raises an important question whether the completeness of the flagellum in each cell temporally correlates with its *qrr* expression and QS response. Therefore, it is important to probe this connection at single-cell level during biofilm formation and host colonization to better understand the correlation between flagellum and QS in a sessile community. As flagellum has been shown to function as a surface sensor in many bacterial species, The unusual link between the flagellum integrity/assembly and QS in *Vibrio* spp. also points to a potential coordination between surface sensing and QS.

Finally, *Vibrio* species are common pathogens that cause diseases in humans and other animals, and they utilize similar QS phosphorelay systems that modulate LuxO phosphorylation and Qrr sRNA levels to regulate virulence gene expression [4, 13, 55–58, 85]. Disruption of the QS pathway in these *Vibrio* species renders the strains non-pathogenic in shrimp, oysters, or mice [4, 5, 13, 86–88]. Thus, the *Vibrio* QS pathways contain several promising targets for drug development to prevent diseases. Therefore, dissecting QS mechanisms in these *Vibrio* species will broadly advance our understanding of the molecular basis of virulence regulation in pathogenic *Vibrio*. Without a thorough understanding the intricacy of QS signaling in these pathogens, ongoing efforts to develop QS-targeted therapeutic strategies are unlikely to succeed.

## Materials and Methods

### Bacterial Strains, Culture conditions

All *V. cholerae* strains used in this study are derivatives of *V. cholerae* O1 El Tor biotype C6706 and E7946. The bacterial strains and plasmids used in this study are listed in Table S1. Bacterial strains were cultured in Luria-Bertani (LB) broth or on LB agar plates at 30°C or 37°C for *V. cholerae* and *E. coli* respectively. Non-cholera *Vibrio* strains were cultured in lysogeny broth marine medium (LM) or in M9 minimal medium supplemented with 0.2% casamino acids and different carbon sources: 20 mM glucose (M9GC) or 1% tryptone w/v (M9TC) at 30 °C. Antibiotics were used at the following final concentrations: streptomycin (sm), 100 µg/mL; kanamycin (kan), 50 µg/mL (25 µg/mL for *E. coli* MG1655); gentamicin, 25 µg/mL; chloramphenicol (cm), 2.5 µg/mL; tetracycline (tet), 10 µg/mL; Spectinomycin (Spec), 100µg/mL; Trimethoprim (Tm), 10 µg/mL and carbenicillin (carb), 100 µg/mL unless otherwise noted.

## Genetic manipulation, transposon excision, and plasmid curing

Construction of *V. cholerae* mutants using allelic exchange with the suicide vector pKAS32 was performed as previously described [64]. Upstream and downstream flanking regions of the target genes were amplified by High-fidelity PCR with Q5 DNA polymerase (New England Biolabs) and cloned into pKAS32 using Gibson assembly. Flagellar transposon insertion mutants were obtained from an order mutant library, a generous gift from Dr. Chris Waters [65]. The transposon insertions with flanking regions were amplified by PCR, gel purified and used for transforming *V. cholerae* recipient (C6706 WT or its isogenic mutants carrying the pMMB-*tfoX-qstR* plasmid, carbenicillin resistant) as previously described [66]. Transformants were selected on LB with kanamycin for transposon insertion. To cure the pMMB-*tfoX-qstR* plasmid after transformation, overnight cultures were diluted 1:1,000 into LB without antibiotic and incubated at 37°C with shaking for 5 hrs to permit plasmid loss. Cultures were then serially diluted and plated onto LB agar with or without carbenicillin and incubated overnight at 37°C. A higher number of colonies on LB plates compared to LB with carbenicillin plates indicated loss of the plasmid from the bacterial population. Colonies from LB plates were screened for carbenicillin sensitivity to confirm plasmid curing.

To excise the kanamycin-resistant transposon, the Flp recombinase plasmid pTL18 [89], (Tet^R^) was first conjugated into the recipient *V. cholerae* transposon mutants. Transconjugants were selected for resistance to both tetracycline and kanamycin. Transconjugants were then grown in LB with tetracycline only at 37 °C with shaking to mid-log phase (OD_600_ 0.4-0.6). Flp recombinase expression was induced by adding isopropyl-β-D-thiogalactopyranoside (IPTG) to a final concentration of 1 mM, followed by incubation for an additional 4–5 hrs at 37°C with shaking. Cultures were serially diluted, plated on LB, and rescreened for sensitivity to both kanamycin and tetracycline. Candidates for which the transposon cassette had been excised leaving the in-frame FRT scar and also lost the pFLP plasmid were confirmed with gene-specific PCR and sequencing.

Non-cholera *Vibrio* mutants were constructed using chitin-independent transformation as described by Simpson et al. [90] and confirmed by Nanopore sequencing.

For plasmid-based complementation or overexpression, the reading frame of the gene of interest was cloned under the control of the *P_tac_* promoter and RBS in pMMB67eh or its gentamycin derivative (pMMBNeo), or pACYC184. Gene expression was induced with 10-100 µM IPTG. Oligonucleotides used in this study are listed in Supplementary Table S2.

## Glucose sensitivity assay

To test for glucose fermentation sensitivity, overnight cultures of different strains were diluted 1000X and inoculated into M9 minimal media with 0.2% casamino acids plus the specified carbon source (0.5% w/v glucose or glycerol). Assays were carried out statically at 30°C. After incubation for 72 hrs, serial dilutions of each sample were plated on LB agar plates to determine viability. Limits of detection are indicated in the figure. Averages and standard errors of mean (SEM) were determined for at least 3 replicates.

## Transformation efficiency assay

The efficiency of chitin-induced transformation of different *V. cholerae* strains was determined as previously described [66]. Briefly, competent cells were prepared by growing different strains in 0.5X Instant Ocean (7 g/L) with chitin slurry (∼0.8 g/mL) at 30°C for overnight. A linear PCR product (*flrC::kan,* ∼500 ng) was then added to the cells and incubated statically at 30°C for 5 hrs. After incubation, LB was added to the cells and grown at 37°C for 2 hrs. Transformants were dislodged from the chitin slurry by vortexing and plated on LB with kanamycin (transformants) and LB (total viable counts) to determine the transformation efficiency (defined as transformants/total viable counts).

## Bacterial two-hybrid assay

Bacterial adenylate cyclase two-hybrid (BACTH) assays were performed using *E. coli* BTH101 cells following the manufacturer’s protocol (Euromedex). The coding sequences of *flgD* and *flrB* were amplified and cloned into both T18 and T25 fusion vectors, generating N- and C-terminal fusions in pUT18, pUT18C, pKT25, and pKNT25 plasmids. The *flrC* gene was cloned into pUT18 and pKNT25 vectors. All constructs were verified by using sequencing and maintained in *E. coli* DH5α.

Different combinations of these T18 or T25 constructs were co-transformed into chemically competent *cyaA*-deficient *E. coli* strain BTH101 cells. Empty vectors were used as negative controls in the co-transformation experiments. A portion of each transformation was plated onto LB agar supplemented with kanamycin (50 µg/mL), carbenicillin (50 µg/mL), 100 µM IPTG, and 40 µg/mL X-gal. Plates were incubated at 30°C for 24 to 48 hrs. Protein-protein interactions were evaluated qualitatively based on the ability of colonies to hydrolyze X-gal, resulting in blue coloration. The color of the colonies was visually inspected. β-galactosidase assay was further performed to quantify *lacZ* gene expression in these transformants using ortho-nitrophenyl-β-galactoside (ONPG) as described [91]. Briefly, colonies grown on selective LB agar plates containing kan, carb, IPTG were scraped and resuspended in water, and cell density was normalized based on OD_600_. A total of 100 µL of normalized cell suspension was added to 900 µL of Z-buffer. Cells were permeabilized by adding 50 µL of 1% (w/v) SDS and 25 µL of chloroform, followed by vortexing and incubation at 30°C for 5 mins. The reaction was initiated by adding 200 µL of ONPG (4 mg/mL). After sufficient yellow color development, the reaction was terminated by adding 500 µL of 1 M Na_2_CO_3_. β-galactosidase activity was calculated in Miller Units [91]. All assays were done in triplicate.

## Luciferase reporter assay

*Vibrio* strains carrying either pBK1003 [13, 82], which harbors the heterologous *luxCDABE* operon fused to the *V. cholerae qrr4* promoter, or a chromosomally integrated P*_qrr4_*–*luxCDABE* fusion at the *lacZ* locus, were streaked onto LB agar supplemented with the appropriate antibiotics. Single colonies were inoculated into LB broth with the corresponding antibiotics and grown overnight (∼16 h) at 30 °C with shaking. Overnight cultures were diluted into fresh LB to equalize cell densities (typically ∼1:1000), and the OD_600_ was adjusted so that all strains started at the same cell density. Non-cholera *Vibrio* strains carrying pBK1003 were grown overnight in LM, washed with M9GC (*V. campbelli*) or M9TC (*V. vulnificus* and *V. parahaemolyticus*) and back-diluted 1:100,000 in 200 µL of the selective minimal media. Diluted cultures (200 µL per well) were dispensed into black 96-well plates, and 50 µL of mineral oil was added to each well to prevent evaporation Plates were incubated at 30 °C for 20 hrs (24 hrs for non-cholera strains), and luminescence and OD_600_ were recorded every 30 min using the BioTek Synergy HT and BioTek Cytation 3 Plate Readers. The LuxO-inhibiting compounds 530A and 530B were synthesized and used as previously described [62]. To induce expression of genes cloned under the P*_tac_* promoter, IPTG was added to the culture medium at final concentrations ranging from 10 to 100 µM.

## Infant mouse model for *in vivo* competition assay

All animal experiments were approved by the Institutional Animal Care and Use Committee at Tufts University School of Medicine (Protocol B2024-26). *Vibrio cholerae* strains were grown overnight (∼16 hrs) at 30°C in LB. The competing strains (one is *lacZ*^+^ and the other is Δ*lacZ*) were mixed at a 1:1 ratio, and the bacterial titer of each inoculum was determined by plating serial dilutions on selective media with X-gal. Approximately 10⁶ CFU of the mixed culture were administered orally to 4- to 7-day-old CD-1 infant mice (Charles River Laboratories). Prior to infection, pups were separated from their mothers 1 hr before inoculation and maintained with restricted access to food and water. Following inoculation, mice were housed at 26°C in the absence of their mothers.

At 24 hrs post-inoculation, infected mice were euthanized by cervical dislocation, and the small intestines were harvested and mechanically homogenized. *V. cholerae* colonization levels were quantified by plating serial dilutions of intestinal homogenates onto LB/Sm/X-Gal agar plates and enumerating colonies after overnight incubation. The competitive index (CI) was calculated as the ratio of each strain in the competing pairs recovered from the intestine, normalized to the ratio present in the input inoculum. A minimum of eight animals was used for each CI determination. Colonization data are presented as individual points per mouse, with the median indicated. For samples in which the strain was below the limit of detection, a value of 1 CFU was assigned at the next lowest dilution at which the other strain was detected (depicted as open symbols in figures).

## Ethics Statement

All animal experiments were performed in accordance with NIH guidelines, the Animal Welfare Act, and US federal law. The infant mouse colonization experimental protocol B2024-26 was approved by Tufts University School of Medicine’s Institutional Animal Care and Use Committee. All animals were housed in a centralized and AAALAC-accredited research animal facility that is fully staffed with trained husbandry, technical, and veterinary personnel in accordance with the regulations of the Comparative Medicine Services at Tufts University School of Medicine.

## Acknowledgement

We thank Dr. Chris Waters for providing the ordered transposon library. We thank Mr. Eliezer Cruz for technical assistance. We thank the members in the Ng and Camilli Labs for their insightful comments and discussions.

## Author Contributions

SSh, SSa, KECV, JCVK, WLN designed research; SSh, SSa, AS, KECV, PW, WLN performed research; SSh, SSa, KECV, JCVK, WLN analyzed data; and SSh, WLN wrote the paper with inputs from KECV and JCVK.

## Notes

### Competing Interest Statement

The authors have declared no competing interest.

### Funding

- National Institutes of Health: R01AI121337, K12GM133314, R35GM124698
- U.S. National Science Foundation: IOS-2525726, BII 2022049

