## supporting information for "Signaling input from the polar flagellum uncouples quorum sensing from cell-density regulation in *Vibrio* species"

**Supplementary Figures Legends:**

**Supplementary Fig. 1** Transcriptional profiles of *qrr*1-4 in WT and Δ*flgD* mutant of *Vibrio cholerae*. Transcription of *qrr*1 (A), *qrr*2 (B), *qrr*3 (C), and *qrr*4 (D) was measured during growth using a bioluminescence reporter (P*qrr*-*luxCDABE*) in WT and Δ*flgD* strains. RLU is relative light units defined as luminescence normalized to $OD_{600}$**.**

**Supplementary Fig. 2** Expression level of VPS-II operon in different *V. cholerae* strains measured with a $P_{vpsL}$-*lux* reporter at different $OD_{600}$.

**Supplementary Fig. 3** Repression of *qrr*4 transcription by FlgD expression in *V. cholerae* flagellar mutants lacking *flgBCD.*

**Supplementary Fig. 4** Schematic representation of both WT and mutated promoter of *qrr*4 fused with a luminescence *lux* reporter. A point mutation (GC to AT) was introduced into the first known LuxO binding site of *qrr*4 promoter (indicated by red color). Not drawn in scale.

### Supplementary Figure 1

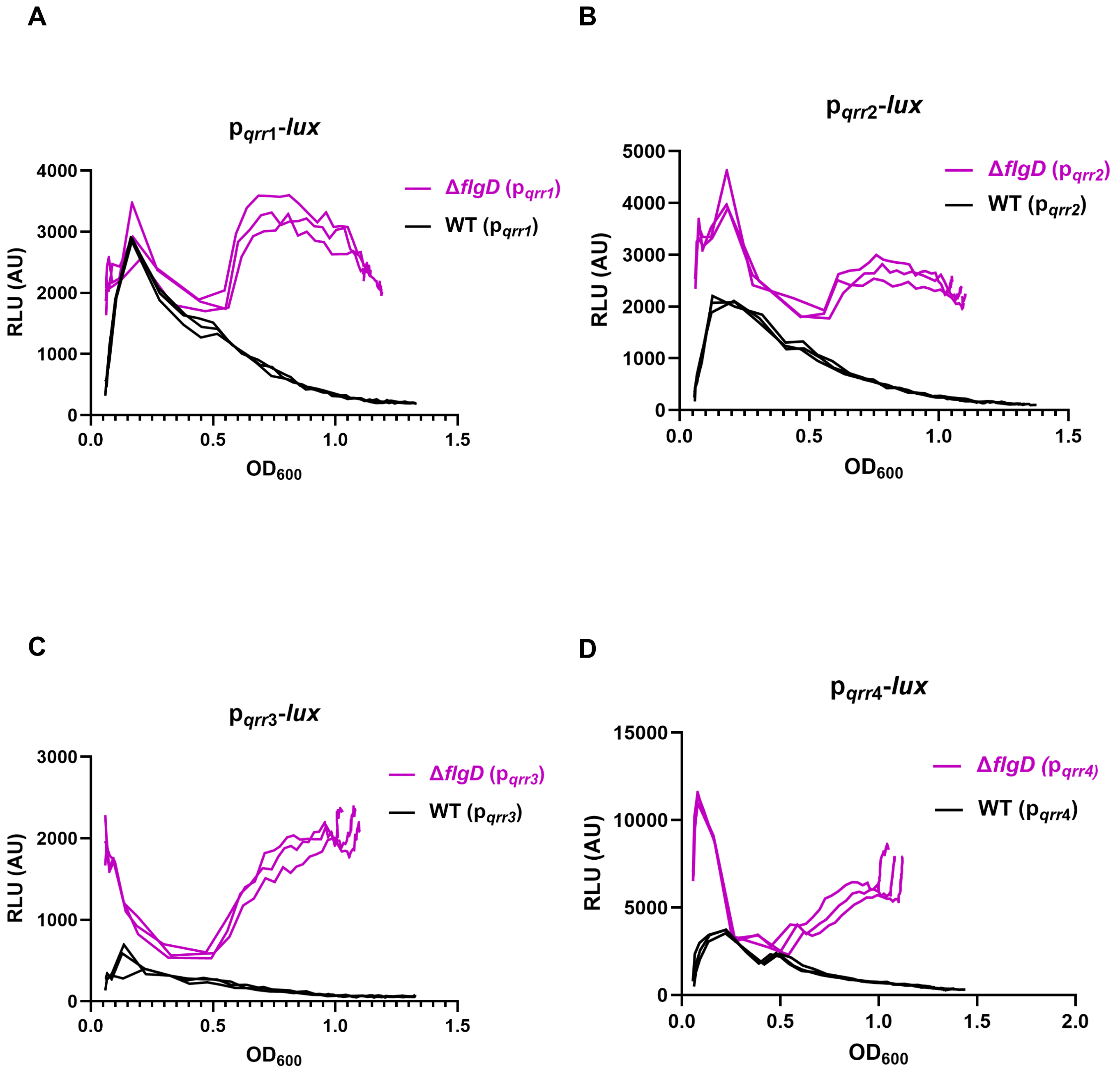

**Supplementary Figure 2**

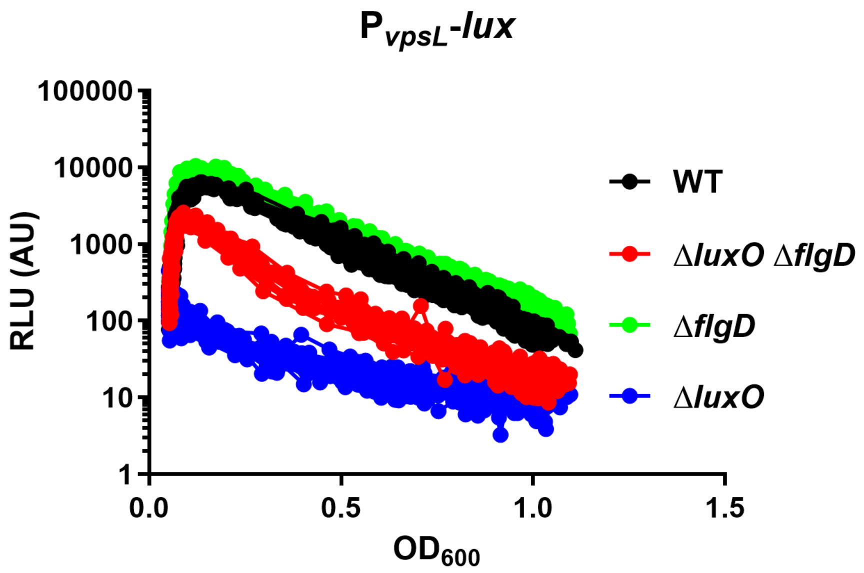

**Supplementary Figure 3**

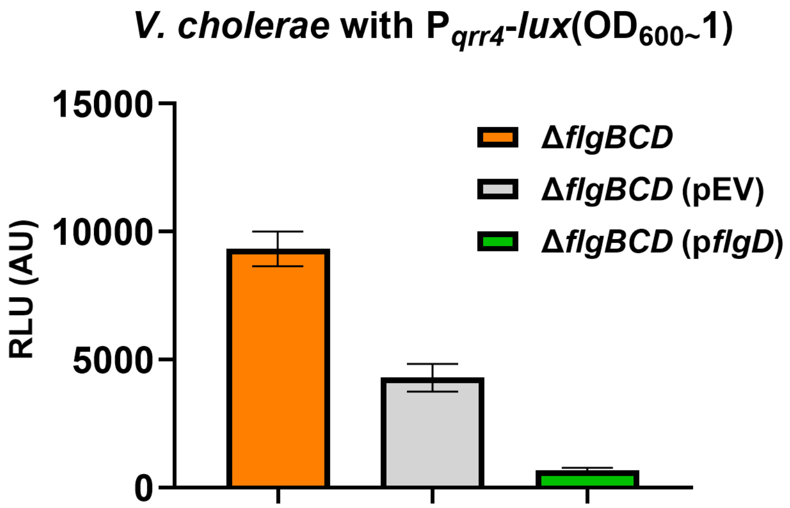

**Supplementary Figure 4**

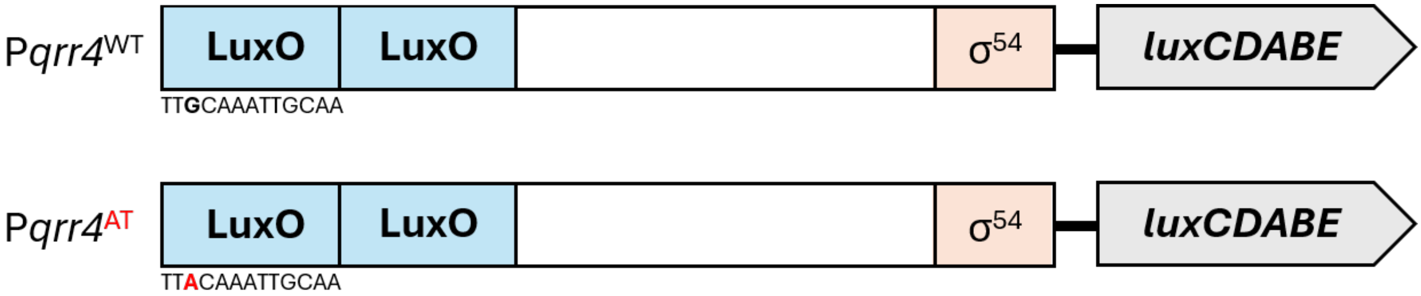

**Supplementary Table 1**

**Bacterial strains and plasmids used in this study**

| **Bacterial Strains/plasmid** | **Description** | **Antibiotic Marker** | **Source** |
|---|---|---|---|
| ***V. cholerae*** | | | |
| WN001 | C6706 | Sm | (1) |
| WN072 | Δ*luxO* | Sm | (2) |
| WN7461 | Δ*flgD* | Sm | This study |
| WN7429 | Δ*luxO* Δ*flgD* | Sm | This study |
| WN6006 | E7946 | Sm | (3) |
| WN5908 | WT *lacZ*::*qrr*4-*lux* | Sm | This study |
| WN8168 | Δ*flgD lacZ*::*qrr*4-lux | Sm | This study |
| WN5910 | Δ*luxO lacZ*::*qrr4*-*lux* | Sm | This study |
| WN7476 | Δ*flgD* Δ*luxO* (pBK1003::P*qrr*4-*lux*) | Sm Cm | This study |
| WN6542 | E7946 (pBK1003 P*qrr*4-*lux*) | Sm Cm | This study |
| WN7243 | E7946 Δ*flgD* (pBK1003::P*qrr*4-*lux* ) | Sm Cm | This study |
| WN7495 | Δ*flgD* Δ*fliA* (pBK1003::*qrr*4- *lux*) | Sm Cm | This study |
| WN6584 | Δ*fliA* VC1807::spec (pBK1003::P*qrr*4-*lux* ) | Sm Cm Spec | This study |
| WN7482 | Δ*flgM* (pBK1003::P*qrr4*-*lux*) | Sm Cm | This study |
| WN6777 | Δ*flgE* VC1807::spec (pBK1003 P*qrr*4-*lux*) | Sm Cm Spec | This study |
| WN6427 | Δ*flrA VC1807*::spec (pBK1003 P*qrr*4-*lux*) | Spec Cm | This study |
| WN6279 | WT *lacZ*::*qrr*4-*lux* Δ*flaACBDE* | Sm | This study |
| WN6781 | Δ*flaACBDE VC1807*::spec *vc2338::kan* (pBK1003 P*qrr4*-*lux)* | Sm Cm Spec Kan | This study |
| WN7466 | Δ*flgD* (pBK1003::P*qrr*3-*lux*) | Cm | This study |
| WN7462 | Δ*flgD* (pBK1003::*qrr1*-*lux*) | Cm | This study |
| WN7464 | Δ*flgD* (pBK1003::*qrr2*-*lux*) | Cm | This study |
| WN6776 | WT *lacZ*::*qrr*4-*lux* Δ*flgE* VC1807::spec | Spec | This study |
| WN6408 | WT *lacZ*::*qrr*4-*lux* Δ*motAB* VC1807::spec | Spec | This study |
| WN8905 | Δ*flgBCD lacZ*::P*qrr4*-*lux* | Sm | This study |
| WN8868 | WT *lacZ*::*qrr4*-*lux flrB*::Tn | Kan | This study |
| WN8864 | WT *lacZ*::*qrr4*-*lux flrC*::Tn | Kan | This study |
| WN8940 | *WT lacZ*::*qrr4*-*lux fliE*::Tn | Kan | This study |
| WN8791 | WT *lacZ*::*qrr4*-*lux flgI*::Tn | Kan | This study |
| WN9006 | *WT lacZ*::*qrr4*-*lux fliP*::Tn | Kan | This study |
| WN8867 | WT *lacZ*::*qrr4*-*lux fliR*::Tn | Kan | This study |
| WN8860 | *WT lacZ*::*qrr4*-*lux flhB*::Tn | Kan | This study |
| WN8859 | WT *lacZ*::*qrr*4-*lux flgB*::Tn | Kan | This study |
| WN8795 | WT *lacZ*::*qrr*4-*lux motX*::Tn | Kan | This study |
| WN8882 | Δ*flgD lacZ*::*qrr4*-*lux flrB*::Tn | Kan | This study |

| | | | |
|---|---|---|---|
| WN8708 | Δ*flgD lacZ*::*qrr4-lux* (pDL1403 pTac *tfox qstR*) *flrC*::Tn | Kan Carb | This study |
| WN8762 | Δ*flgD lacZ*::*qrr4-lux* (pDL1403 pTac *tfox qstR*) *fliE*::Tn | Kan Carb | This study |
| WN8783 | Δ*flgD lacZ*::*qrr4-lux* (pDL1403 pTac *tfox qstR*) *flgI::Tn* | Kan Carb | This study |
| WN8886 | Δ*flgD lacZ*::*qrr4-lux fliP*::Tn | Kan | This study |
| WN8710 | Δ*flgD lacZ*::*qrr4-lux* (pDL1403 pTac *tfox qstR*) *fliR*::Tn | Kan Carb | This study |
| WN8884 | Δ*flgD lacZ*::*qrr4-lux flhB*::Tn | Kan | This study |
| WN8704 | Δ*flgD lacZ*::*qrr4-lux* (pDL1403 pTac *tfox qstR*) *motX*::Tn | Kan Carb | This study |
| WN8761 | Δ*flgD lacZ*::*qrr4-lux* (pDL1403 pTac *tfox qstR*) *motY*::Tn | Kan Carb | This study |
| WN8858 | WT *lacZ*::*qrr*4-*lux flrB*::FRT | Sm | This study |
| WN9007 | WT *lacZ*::*qrr*4-*lux flrC*::FRT | Sm | This study |
| WN8996 | WT *lacZ*::*qrr*4-*lux fliE*::FRT | Sm | This study |
| WN9291 | WT *lacZ*::*qrr*4-*lux flgI*::FRT | Sm | This study |
| WN9015 | WT *lacZ*::*qrr*4-*lux fliP*::FRT | Sm | This study |
| WN9000 | WT *lacZ*::*qrr*4-*lux fliR*::FRT | Sm | This study |
| WN9014 | WT *lacZ*::*qrr*4-*lux flhB*::FRT | Sm | This study |
| WN9061 | WT *lacZ*::*qrr*4-*lux motX*::FRT | Sm | This study |
| WN9063 | Δ*flgD lacZ*::*qrr4-lux flrB*::FRT | Sm | This study |
| WN8999 | Δ*flgD lacZ*::*qrr4-lux flrC*::FRT | Sm | This study |
| WN9059 | Δ*flgD lacZ*::*qrr4-lux fliE*::FRT | Sm | This study |
| WN9290 | Δ*flgD lacZ*::*qrr4-lux flgI*::FRT | Sm | This study |
| WN9001 | Δ*flgD lacZ*::P*qrr4-lux fliP*::FRT | Sm | This study |
| WN9016 | Δ*flgD lacZ*::P*qrr4-lux fliR*::FRT | Sm | This study |
| WN9002 | Δ*flgD lacZ*::P*qrr4-lux flhB*::FRT | Sm | This study |
| WN8997 | Δ*flgD lacZ*::P*qrr4-lux motX*::FRT | Sm | This study |
| WN8998 | Δ*flgD lacZ*::P*qrr4-lux motY*::FRT | Sm | This study |
| WN9047 | WT *lacZ*::P*qrr4-lux:flrB*::FRT (pMMB *flrC*$^{M114I}$) | Gm | This study |
| WN9050 | WT *lacZ*::P*qrr4-lux flrB*::FRT (pMMB EV) | Gm | This study |
| WN9049 | *flrC*::FRT *lacZ*::P*qrr4-lux* (pMMB *flrC*$^{M114I}$) | Gm | This study |
| WN9052 | *flrC*::FRT *lacZ*::P*qrr4-lux* (pMMB EV) | Gm | This study |
| WN9054 | Δ*flgD lacZ*::P*qrr4-lux flrC*::FRT(pMMB *flrC*$^{M114I}$) | Gm | This study |
| WN9046 | Δ*flgD lacZ*::P*qrr4-lux flrC*::FRT (pMMB EV) | Gm | This study |
| WN8899 | Δ*flgD lacZ*::*qrr4-lux fliP*::Tn (pMMBgm::pTac *flgD*+) | Kan Gm | This study |
| WN8931 | Δ*flgD lacZ*::*qrr4 fliP*::Tn (pMMB EV) | Kan Gm | This study |
| WN8901 | Δ*flgD lacZ*::*qrr4-lux flhB*::Tn (pMMBgm::pTac *flgD*+) | Kan Gm | This study |

| | | | |
|---|---|---|---|
| WN8929 | Δ*flgD lacZ*::*qrr4 flhB*::Tn (pMMB EV) | Kan Gm | This study |
| WN8966 | *ΔflgBCD lacZ*::P*qrr4-lux* (pMMB EV) | Gm | This study |
| WN8942 | *ΔflgBCD lacZ*::P*qrr4-lux* (pMMB *flgD*) | Gm | This study |
| WN9082 | C6706 (pBk1003_P*qrr*$^{AT}$-*lux*) | Cm | This study |
| WN9100 | C6706 (pBk1003_P*qrr*$^{AT}$-*lux*+pMMB *flrC*$^{M114I}$) | Cm Gm | This study |
| WN9104 | C6706 (pBk1003_P*qrr*$^{AT}$-*lux*+pMMB EV) | Cm Gm | This study |
| WN9080 | C6706 Δ*flgD*(pBk1003_P*qrr*-lux*) | Cm | This study |
| WN9102 | C6706 Δ*flgD*(pBk1003_P*qrr*-lux*+pMMB *flrC*$^{M114I}$) | Cm Gm | This study |
| WN9106 | C6706 Δ*flgD*(pBk1003_P*qrr*$^{AT}$-*lux*+pMMB EV) | Cm Gm | This study |
| ***E. coli*** | | | |
| WN5870 | S17/pKAS *lac*::P*qrr4-lux* | Kan | This study |
| WN7472 | S17λpir (pkas32 *ΔfliA*) | Am | This study |
| WN7229 | S17λpir (pKAS32 *ΔflgM*) | Am | This study |
| WN6351 | S17λpir DAP- / pSLS54 ($P_{qrr3}$-*lux*) | DAP$^{-}$, Cm | (4) |
| WN7318 | S17λpir (pBK1003*qrr*1-*lux*) | Cm | This study |
| WN7319 | S17λpir pBK1003*qrr*2-*lux*) | Cm | This study |
| WN7100 | S17λpir (pKAS32 Δ*flgD*) | Am | This study |
| WN042 | WT MG1655 | - | Lab collection |
| WN8961 | MG1655 *flgD*::kan (pBK1003*qrr*4-*lux* pMMB *flrC*M114I) | Kan Cm Gm | This study |
| WN8630 | MG1655 *flgD*::kan (pMMB EV pBK1003 pBK1003*qrr*4-*lux*) | Kan Cm Gm | This study |
| WN8963 | MG1655 WT (pBK1003 *qrr*4-*lux* pMMB *flrC*M114I) | Gm Cm | This study |
| WN8957 | MG1655 WT (pMMB EV+pBK1003 *qrr*4-*lux*) | Gm Cm | This study |
| WN8980 | S17λpir (pMMB *flrBC*) | Kan | This study |
| WN8907 | S17λpir (pMMB *flrC*M114I) | Gm | This study |
| WN8908 | S17λpir (pMMB EV) | Gm | This study |
| WN8898 | Dh5aλpir (pMMB *flrC*M114I) | Gm | This study |
| WN8897 | S17λpir (pMMB *flgD*) | Gm | This study |
| WN8892 | Dh5aλpir (pMMB *flgD*) | Gm | This study |
| WN8891 | Dh5aλpir (pkAS32 Δ*flgBCD*) | Kan | This study |
| WN8896 | S17λpir (pkAS32 Δ*flgBCD*) | Kan | This study |
| WN9071 | S17λpir (pBk1003 P*qrr*4$^{AT}$-*lux*) | Cm | This study |
| WN9088 | MG1655 WT (pBk1003 P*qrr*4-*lux**+pMMB *flrC*M114I) | Cm Gm | This study |
| WN9090 | MG1655 flgD::kan (pBk1003 P*qrr*4$^{AT}$-*lux*+pMMB *flrC*M114I) | Kan Cm Gm | This study |
| WN9096 | MG1655 WT (pBk1003 P*qrr*4$^{AT}$-*lux*+pmmB EV) | Cm Gm | This study |

| | | | |
|---|---|---|---|
| WN9098 | MG1655 flgD::kan (pBk1003 P*qrr*4$^{AT}$-*lux*+pMMB EV) | Kan Cm Gm | This study |
| WN9273 | MG1655 WT (pACYC184 *flgD* his6) | Tet | This study |
| WN9295 | MG1655 WT (pACYC184 EV+pMMB *flrBC*+pBK1003 *qrr*4-*lux*) | Kan Cm Tet | This study |
| WN9189 | DH5αλpir (pKNT25 *flrC*) | Kan | This study |
| WN9181 | DH5αλpir (pUT18 *flrC*) | Carb | This study |
| WN9188 | DH5αλpir( pUT18 *flgD* ) | Carb | This study |
| WN9239 | Dh5αλpir (pKT25C *flgD*) | Kan | This study |
| WN9241 | Dh5αλpir (pUT18C *flgD*) | Carb | This study |
| WN9201 | DH5αλpir (pUT18 *flrB*) | Carb | This study |
| WN9204 | DH5αλpir (pUT18 *flrC*) | Carb | This study |
| WN9207 | Dh5αλpir (pKNT25 *flgD*) | Kan | This study |
| WN9208 | Dh5αλpir (pKNT25 *flrB*) | Kan | This study |
| WN9301 | Dh5αλpir (pUT18C *flrB*) | Carb | This study |
| WN9302 | Dh5αλpir (pKT25 *flrB*) | Kan | This study |
| WN8343 | BTH101 | Sm | Euromede x |
| WN9273 | MG1655 WT (pACYC184 *flgD*-his6) | Tet | This study |
| WN9295 | MG1655 WT (pACYC184 EV+pMMB *flrBC*+pBK1003 P*qrr*4-*lux*) | Kan Cm Tet | This study |
| WN9296 | MG1655 WT (pACYC184 *flgD*-his6+pMMB *flrBC*+pBK1003 P*qrr*4-*lux*) | Kan Cm Tet | This study |
| WN9311 | MG1655 (pACYC184 FlgD his6+pmmBgm FlrCM114I+pBK1003P*qrr*4-*lux*) | Gm Cm Tet | This study |
| WN9349 | MG1655 (pACYC184 EV+pMMBgmFlrCM114I+pBK1003P*qrr4* -*lux* | Gm Cm Tet | This study |
| **Non cholera strains** | | | |
| DS40M4 | Wild-type *V. campbelli* | | (5) |
| CAS197 | DS40M4, Δ*luxO,* Δ*luxB*::$spec^R$ | Spec | (6) |
| BDP109 | DS40M4, pMMB67EH-tfox-kanR, Δ*luxB*::$spec^R$, Δ*flgD* | Kan Spec | (7) |
| ARK001 | DS40M4, Δ*luxO,* Δ*flgD,* Δ*luxB*::$tm^R$, pMMB67EH-tfox-kanR | Tm Kan | This study |
| ATCC 27562 | Wild-type *V. vulnificus* | | ATCC |
| CAS-vv022 | ATCC 27562, pCS32 | | (8) |
| KECV0011 | ATCC 27562, Δ*flgD::*$tm^R$, pCS32 | Tm | This study |
| RIMD2210633 | Wild-type *V. parahaemolyticus* | | ATCC |
| CAS-V01 | RIMD2210633, pMMB67EH-tfoX-kanR | Kan | (8) |
| KECV0012 | RIMD2210633, Δ*flgD::*$tm^R$, pMMB67EH-tfoX-kanR | Tm Kan | This study |

Sm, streptomycin; Kan, kanamycin; Spec, spectinomycin; Tet, tetracycline; Gm, gentamicin; Cm, chloramphenicol; Carb, Carbenicillin; Tm, trimethoprim; Amp, ampicillin; DAP, diaminopimelic acid.

**Supplementary Table 2:**

**Primers used in this study**

| Name | Sequence | Target |
|---|---|---|
| WNTP0540 | GCGGCCGCCTGCAGCTGG | Gibson Primer for pKAS32 |
| WNTP0541 | CATATGCATCCTAGGCCTATTAATATTCCGGAGTATACGTAGCC | |
| WNTP2462 | GGTAAACATGGCTATTTCTTTTGACAG | amplified *flgB* for Tn transformation |
| WNTP2463 | CTCATCTATCTACTCCCCTTTAAGC | |
| WNTP2464 | GGCTACCCGTGATATTGCTG | TnFGL3_rev |
| WNTP2487 | CTGATTTGGCAGAATCCGAC | amplify *flhB* for Tn transformation |
| WNTP2488 | CTCATTTAGTAACGCATATCCGG | |
| WNTP2489 | GAGTGCGGGTATGAAGCTAC | amplify *motX* for Tn transformation |
| WNTP2490 | GACAGCAATTACCAGAACGTTTC | |
| WNTP2491 | CGCCTCAATTAAATGAATAAATGGG | amplify *motY* for Tn transformation |
| WNTP2492 | TCCACCAATCACACCTGAG | |
| WNTP2504 | GCCATCTTCACTCTCCAGTCTC | amplify *flrB* for Tn transformation |
| WNTP2505 | GAGTGAATGATGAGCAGCGC | |
| WNTP2506 | GTATGCCATGCCCTCAACC | amplify *flrC* for Tn transformation |
| WNTP2507 | GACTGGAGAGTGAAGATGGCTC | |
| WNTP2510 | CTTGTCCGTCGGATTCTGC | amplify *fliR* for Tn transformation |
| WNTP2511 | GCAGGTTCTGTACTAGGAGTCC | |
| WNTP2520 | catttttATGGCCGGAGTCAATAATGTTGG | amplify *flgD* for pmmBgm |
| WNTP2521 | gcagttaATGGTGGTGATGGTGATGTCACGCTTTGCCTACTTCAAGTAC | |
| WNTP2522 | TTGACTCCGGCCATaaaaatgtatcctaaagtcgactctagag | pmmBgm_*flgD* (backbone) |
| WNTP2523 | CCATCACCACCATtaactgcaggcatgcaagc | |
| WNTP2530 | CTCTGGTGTCATGTGCCC | |

| | | |
|---|---|---|
| WNTP2 531 | CTAACCAGTTGATGCAGCTACTG | amplify *fliP* for Tn transformation |
| WNTP2 532 | CGCAGTAGCTAATTTAAGGTAGGAGTTA | amplify *flgG* for Tn transformation |
| WNTP2 533 | GCGGCCATCTCTTAGTTAGC | |
| WNTP2 536 | TCATCGGCCACAGTCTATTACC | amplify *fliE* for Tn transformation |
| WNTP2 537 | GTCAAACAATTGGCATCAGAGGTAG | |
| WNTP2 545 | GATTGAGGACGAGAAAGAGTGC | amplify *flgJ* for Tn transformation |
| WNTP2 546 | GCGATTGAAGGCCAACTGATC | |
| WNTP2 547 | GCGTAACTTGTCGAGTCCG | amplify *flgI* for Tn transformation |
| WNTP2 548 | GCATAGTGAACGGTGAGGC | |
| WNTP2 563 | caggcggccgcTTAACGAATCTGCAGGATATTTTGTTGC | amplify of *flgBCD* deletion fragment for pKAS-kan |
| WNTP2 564 | GAAGAGGTAAACATGCTTGAAGTAGGCAAAGCGTGA | |
| WNTP2 565 | CTTCAAGCATGTTTACCTCTTCGATAAGAACTGACC | |
| WNTP2 559 | taggcctaggatgcatatgATGACTGCTATAACGATTAGCGACC | |
| WNTP2 591 | ctggtgcgtaacggctgaTa | Screening *flrC*$^{M114I}$ mutant_R |
| WNTP2 580 | agagtcgactttaggatacattttCCTATTGCAGTCACGGATCAGTG | clone of *flrC*$^{M114I}$ into pmmBgm_fragment1 |
| WNTP2 568 | AACGGCTGACAATATTCAGCA | |
| WNTP2 167 | aaaaatgtatcctaaagtcgactctagaggattcc | pmmBgm gibson linearization |
| WNTP2 581 | taactgcaggcatgcaagcttg | |
| WNTP2 582 | cttgcatgcctgcagttaTCAACCGGGGATATCAATCCCTG | clone of *flrC*$^{M114I}$ into pmmBgm_fragment2 |
| WNTP2 569 | GCTGCTGAATATTGTCAGCCGTTAC | |
| WNTP1 569 | aaggcaaataatctcacctggaaaccatagtcaatatattg | *flrA* deletion |
| WNTP1 570 | atatattgactatggtttccaggtgagattatttgcctttatt | |
| WNTP1 571 | GACAGATCTGCGCGCGATCgtgagcgagtggtgaaaaag | |
| WNTP1 572 | aacggctgacatgggaattcgtggttaatgtggaagcagaa | |

| | | |
|---|---|---|
| WNTP1593 | aattgcaatgatcatcatgtttcgcctgcagcagaaag | *motAB* deletion |
| WNTP1594 | attttGagtgacacaggaacaaatcgcactttggccgatg | |
| WNTP1633 | gcctgtactttagtcaatgatg | *motY* deletion |
| WNTP1631 | gaaagtctacatcgtctctgc | |
| WNTP1632 | cggtctctacatcgataacc | *motY* sequencing |
| WNTP1634 | ccaagttttggcgcaccatg | |
| WNTP1641 | cgaaatcatcaatcttgaattag | deletion of *flaAC* |
| WNTP1639 | gaatgatttccatgagacggt | |
| WNTP1640 | caaaagagaccaaagctgacc | sequencing of *flaCD* |
| WNTP1642 | caggcaataaaaaacccagctg | |
| WNTP1646 | ccttgctgcattgccgcc | Deletion of *flaBDE* |
| WNTP1643 | cgcttaagaaaaaagagcgc | |
| WNTP1644 | caccgataaaagaacaagtaag | Sequencing of *flaDE* |
| WNTP1645 | tgaaaaaaagcctgaaatttctc | |
| WNTP1681 | gaaagctgagaaacgtgcg | sequencing of *flaBDE* |
| WNTP1682 | gatcctgttcacgggcaag | |
| WNTP1737 | tatggcaggtattgataacggcggcttctaacctaacgtg | *flgM* deletion |
| WNTP1738 | gcacctgtttgcgtgccac | |
| WNTP1739 | gcattactgctcgtacgcc | |
| WNTP1740 | acgttaggttagaagccgccgttatcaatacctgccataaatg | |
| WNTP1725 | gtggctaaattggcggctc | *fliA* deletion |
| WNTP1726 | agtgatatttagtcattccggtattgatcgtatgtaagcgc | |
| WNTP1727 | cgcttacatacgatcaataccggaatgactaaatatcactgat | |
| WNTP1728 | gttgcagtggttcacgctc | |
| WNTP1753 | ggtcactgggggtaaaggt | *cheY*-3 deletion |

| | | |
|---|---|---|
| WNTP1754 | Gcagttgatcaagtagcttttc | |
| WNTP1755 | ccgtttgaagagggggaag | sequencing of *cheY*-3 |
| WNTP1756 | ggaatttcatccgtcgcgat | |
| WNTP1718 | cacgctttgcctacttcaaggactccggccatacgttaac | *flgD* deletion |
| WNTP1719 | agttaacgtatggccggagtccttgaagtaggcaaagcgtg | |
| WNTP1805 | aggaggAAGCTTaacgtgctgatctatttctcg | |
| WNTP1806 | tcctccGCTAGCcttgctcgttcgcaacacg | |
| WNTP1745 | gcgaaagatacttacaaagct | sequencing of *flgD* |
| WNTP1746 | ggaactaagcgatctttcgc | |
| WNTP1815 | ccgcaaaacttaatggaaagtt | *flgE* deletion |
| WNTP1816 | cgccaaattaaacagcgagc | |
| WNTP1817 | taacgaatctggacatatgacatgagtaattctc | |
| WNTP1818 | aattactcatgtcatatgtccagattcgttaatcctcttcaa | |
| WNTP1823 | gaacctttgacgtcggtgcc | sequencing of *flgE* |
| WNTP1824 | ccgtaagcctgcatagaac | |
| WNTP2003 | ccttaaggctctctggcttagcggccgcggcatccgaatccctgac | *flgD* complementation into pKAS |
| WNTP2004 | acattattgactccggccatgtttacctcttcgataagaac | |
| WNTP2005 | atcgaagaggtaaacatggccggagtcaataatg | |
| WNTP2006 | actcctcggcttgagggatgggtacctcacgctttgcctacttcaag | |
| WNTP2060 | cctgactgggctgacgg | Sequencing of *flgD* |
| WNTP2061 | tcacgctttgcctacttcaag | |
| WNTP0782 | actcctcggcttgagggatgATTCCCATGGACATAGCG | clone of P*qrr*4-*lux* into pBK1003 |
| WNTP0783 | ttaattgtcaacagggtaccTCAACTATCAAACGCTTCG | |
| WNTP0803 | ccttaaggctctctggcttaTCAACTATCAAACGCTTCG | Gibson primer to insert *qrr*4-*lux* to pKAS-*lac* |
| WNTP0804 | TAAGCCAGAGAGCCTTAAGGCTCTCTTTTTTGTG | |

| | | |
|---|---|---|
| WNTP1959 | cggccgctctagaACTAGTCACCGACGCCGTGTCTTC | *qrr*1 fwd pBBRlux SpeI gibson |
| WNTP1960 | gcggccgcaactagaGGATCCCAGTAGTAATCAAGCACATATC | *qrr*1 rev pBBRlux BamHI gibson |
| WNTP1961 | cggccgctctagaACTAGTGAATAAGGCCGTCATTATACC | *qrr*2 fwd pBBRlux SpeI gibson |
| WNTP1962 | gcggccgcaactagaGGATCCCACCTAAACTAATGCACGAAG | *qrr*2 rev pBBRlux BamHI gibson |
| WNTP2661 | tgattacgccaagcttgcatatggccggagtcaataatg | pKNT25-*flgD* fwd |
| WNTP2662 | tctagagtcgacctgcaggccgctttgcctacttcaag | |
| WNTP2663 | caagcttgcatgcctgcaggatggccggagtcaataatg | pUT18-*flgD* |
| WNTP2664 | cggggatcctctagagtcgacgctttgcctacttcaag | |
| WNTP2665 | tgattacgccaagcttgcatatgatgagcagcgcagtg | pKNT25-*flrB* |
| WNTP2666 | tctagagtcgacctgcaggcctctccagtctctgagtttg | |
| WNTP2667 | caagcttgcatgcctgcaggatgatgagcagcgcagtg | pUT18-*flrB* |
| WNTP2668 | cggggatcctctagagtcgactctccagtctctgagtttg | |
| WNTP2669 | tgattacgccaagcttgcatatgcagagtttagcgaaac | pKNT25-*flrC* |
| WNTP2670 | tctagagtcgacctgcaggcgcgttgcatgttgtatttg | |
| WNTP2671 | caagcttgcatgcctgcaggatgcagagtttagcgaaac | pUT18-*flrC* |
| WNTP2672 | cggggatcctctagagtcgagcgttgcatgttgtatttg | |
| WNTP2678 | GGTCGACTCTAGAGGATCCCatggccggagtcaataatgttgg | *flgD*-pUT18C |
| WNTP2679 | CGAGCTCGGTACCCGcgctttgcctacttcaagtacttcag | |
| WNTP2680 | AGGGTCGACTCTAGAGGATatggccggagtcaataatgttgg | *flgD*-pKT25 |
| WNTP2681 | ACTTAGGTACCCGGGGcgctttgcctacttcaagtacttcag | |
| WNTP2698 | GGTCGACTCTAGAGGATCCCATGATGAGCAGCGCAGTGC | *flrB*-pUT18C |
| WNTP2699 | CGAGCTCGGTACCCGCTCTCCAGTCTCTGAGTTTGAGCTATC | |
| WNTP2700 | AGGGTCGACTCTAGAGGATATGATGAGCAGCGCAGTGC | pKT18-*flrB* |

| | | |
|---|---|---|
| WNTP2 701 | gTTACTTAGGTACCCGGGGCTCTCCAGTCTCTGAG TTTGAGCTATC | |
| WNTP2 674 | ttgacaattaatcatcggctcgtataatgtgtggaattgtgagcggataaca aAATTAACTGAAAGGAGGTTAAAATGGCCGGAGTC AATAATGTTGG | clone of *flgD*_his6 into pACYC184 |
| WNTP2 675 | cggaattccTCAATGGTGGTGATGGTGATGCGCTTTG CCTACTTCAAGTACTTCA | |
| WNTP2 676 | tacgagccgatgattaattgtcaaggatgagcattcatcaggcgg | gibson linearization of pACYC184(*flgD*-his6) |
| WNTP2 677 | CACCACCATTGAggaattccgtatggcaatgaaagacg | |
| WNTP2 636 | ctagagtcgactttaggatacatttttATGATGAGCAGCGCAGT GC | clone of flrBC into pMMBneo |
| WNTP2 630 | gcttgcatgcctgcagTCAACCGGGGATATCAATCCC | |
| WNTP 2617 | aaaaatgtatcctaaagtcgactctagag | gibson linearization of pMMBneo |
| WNTP 2097 | aggtcatgagccagacccag | |
| WNTP 2616 | ctgcaggcatgcaagcttg | |
| WNTP 2096 | tgggtctggctcatgacctg | |
| WNTP2 600 | GACGAAATGCGATTTCTATTaCAAATTGCAAAATCA A | introduction of point mutation in first LuxO binding site on *qrr*4 promoter |
| WNTP2 601 | CACCTCGACCTGAATGGAAGCC | |
| WNTP2 602 | GATTTTGCAATTTGtAATAGAAATCGCATTTCGTC | |
| WNTP2 603 | CATCCAACGTTGATGAATATCCACCTC | |
| WNTP2 604 | CCATTCAGGTCGAGGTGGC | Linearization of pBk1003-*qrr-lux* |
| WNTP2 605 | GGTGGATATTCATCAACGTTGGATGATTATTGAG | |
| KECV00 55 | tcatgcttgccagtggtagagtatc | Δ*flgD* RIMD2210633 F1 |
| KECV00 66 | agacttctcaggtcgacggatccccggaatggccatacgctacctccttat cc | Δ*flgD* RIMD2210633 R1 |
| KECV00 67 | gcaacggattcgaagcagctccagcctacataatccataacatgctagat taggag | Δ*flgD* RIMD2210633 F2 |
| KECV00 58 | cctgttggcaagtaaaggttctgac | Δ*flgD* RIMD2210633 R2 |
| KECV00 61 | ccataccgtgtcggtgattgac | Δ*flgD* ATCC 27562 F1 |

| | | |
|---|---|---|
| KECV0068 | agacttctcaggtcgacggatccccggaatggccatacgctacctccttatcc | Δ*flgD* ATCC 27562 R1 |
| KECV0069 | gcaacggattcgaagcagctccagcctacagcgtaattacgctggctagattcag | Δ*flgD* ATCC 27562 F2 |
| KECV0064 | gttacgttcacgttggatgtttctagc | Δ*flgD* ATCC 27562 R2 |
| CAS0074 | gtccgtcgctctaccaactgagc | Δ*luxO* DS40M4 F1 |
| CAS0297 | gctaattcagtttaagcggccattaccattagtagataacgagac | Δ*luxO* DS40M4 R1 |
| CAS0298 | atggccgcttaaactgaattagcgtatgaatacggacgtattaaatcagc | Δ*luxO* DS40M4 F2 |
| CAS0079 | gtgcttctggcgtgctgtcacg | Δ*luxO* DS40M4 R2 |
| CAS0148 | cgtgctcaagtcttcactgatgatg | Δ*luxB* DS40M4 F1 |
| CAS0149 | gtcgacggatccccggaatgatgacttgatcagaagaacgctttga | Δ*luxB* DS40M4 R1 |
| CAS0150 | gaagcagctccagcctacacactcgtaacgtttaaacgatgctgag | Δ*luxB* DS40M4 F2 |
| CAS0151 | ggtgaatggccacaaggtacct | Δ*luxB* DS40M4 R2 |
| ABD0123 | attccggggatccgtcgac | Ampflify AbR ($spec^R$, $tm^R$) F |
| ABD0124 | tgtaggctggagctgcttc | Ampflify AbR ($spec^R$, $tm^R$) R |
